# *Arabidopsis thaliana* rosette growth habit is a photomorphogenic trait controlled by the TALE homeodomain protein ATH1 and involves TOR kinase

**DOI:** 10.1101/2022.05.13.491803

**Authors:** Shahram Shokrian Hajibehzad, Savani S. Silva, Niels Peeters, Evelien Stouten, Guido Buijs, Sjef Smeekens, Marcel Proveniers

**Affiliations:** Molecular Plant Physiology, Department of Biology, Science4Life, Utrecht University, Padualaan 8, Utrecht, 3584 CH, The Netherlands

**Author notes:** Correspondence: Marcel Proveniers.

**Keywords:** *ARABIDOPSIS THALIANA* HOMEOBOX 1 (ATH1), rosette growth habit, photomorphogenesis, TOR kinase, meristem activity

## Abstract

Here, we demonstrate that Arabidopsis rosette habit is a *bona fide* photomorphogenic trait controlled by the homeodomain protein ATH1. In light, *ATH1* expression at the SAM is induced by broad wavelengths, mediated through multiple photoreceptors, and requires inactivation of COP1 and PIF photomorphogenesis inhibitors. Such induced *ATH1* prevents elongation of rosette internodes by maintaining the rib zone area of the SAM in an inactive state. In the absence of light, Arabidopsis plants cannot complete seedling establishment after germination due to inactivity of the shoot apical meristem (SAM). Light requirement for SAM activation can be overcome by availability to the meristem of metabolizable sugars, such as sucrose. However, under these conditions plants fail to establish a typical compact rosette and display a caulescent growth habit. We show that this is due to insufficient expression of *ATH1* at the SAM. ATH1 induction restores rosette habit in dark-grown plants through inhibition of *PIF* gene expression. Together, this suggests that a SAM-specific, double-negative ATH1-PIF feedback loop is at the basis of Arabidopsis rosette habit. Induction of *ATH1* expression and restoration of rosette habit in darkness also occurs at increased levels of sucrose. Both sugar and light signals that induce *ATH1* are mediated by TOR kinase. Overall, these results support a fundamental role for ATH1 in Arabidopsis rosette habit and further strengthen a role for TOR kinase as a central hub for integration of energy and light signals controlling organogenesis at the SAM.

## Introduction

Plants, as sessile organisms, are equipped with sophisticated mechanisms to sense the environment and to adapt their growth and development accordingly. Being photoautotrophs, plants are especially attuned to the light environment. This is well illustrated by the dramatic differences in appearance between light- and dark-grown seedlings. In Arabidopsis, dark-grown seedlings have a typical etiolated phenotype, characterized by an elongated hypocotyl, formation of an apical hook, closed and unexpanded cotyledons, and an arrested shoot apical meristem (SAM). Exposure of seedlings to light results in inhibition of hypocotyl elongation, opening of the apical hook, opening and expansion of cotyledons, and SAM activation (Chen and Chory, 2011; Arsovski et al., 2012; Pfeiffer et al., 2016; Mohammed et al., 2017; Janocha et al., 2021). The active SAM gives rise to the aerial portion of the plant via the formation of organ primordia at its flanks. During the vegetative growth phase, leaf primordia arise in a spiral phyllotaxy to form a compact, basal rosette in which internode elongation remains arrested. In the absence of light, SAM activity can be induced by exposing the SAM to metabolizable sugar, such as sucrose, glucose, or fructose (Araki and Komeda, 1993; Roldán et al., 1999). Both light- and sugar-mediated SAM activation involve TARGET OF RAPAMYCIN (TOR) kinase, a central component in energy sensing, such that it promotes SAM activity in favorable conditions (Pfeiffer et al., 2016; Li et al., 2017; Mohammed et al., 2017; Janocha et al., 2021). It has been proposed that light, via photoreceptor signaling through CONSTITUTIVE PHOTOMORPHOGENIC1 (COP1), plays a permissive role toward energy signaling in the SAM, possibly by controlling sugar import into the meristem (Mohammed et al., 2017). This might explain why direct access of the SAM to metabolizable sugar can activate the meristem in the absence of light.

Sugar-induced dark morphogenesis of Arabidopsis plants follows the same developmental phases as light-grown plants. However, in sugar-induced, dark-grown plants stem elongation is not inhibited during vegetative development contrary to light-grown plants. Consequently, such dark-grown plants no longer display a rosette habit and elongated internodes are present between adjacent ‘rosette’ leaves (Roldán et al., 1999; Mohammed et al., 2017). A similar loss of rosette habit has been observed in light-grown Arabidopsis plants lacking functional phytochrome (phy) and/or cryptochrome (CRY) photoreceptors. Control of rosette internode elongation in response to a low red to far-red (R:FR) ratio of light or end-of-day FR light (EOD-FR) is mediated by phyA, phyB, phyD, and phyE in a functionally redundant manner (Devlin et al., 1996; Whitelam and Devlin, 1997; Devlin et al., 1998; Whitelam et al., 1998; Devlin et al., 1999; Roldán et al., 1999; Mazzella et al., 2000; Devlin et al., 2003; Franklin et al., 2003a). In addition, ambient temperature has been reported to modulate the light-regulation of Arabidopsis rosette habit. At elevated ambient temperature, phyB and CRY1 redundantly suppress internode elongation during vegetative development (Mazzella et al., 2000). A compact rosette habit thus is a *bona fide* photomorphogenic trait in Arabidopsis. However, despite numerous observations and the economic importance of rosette habit in vegetable crops, this aspect of photomorphogenic development has been paid little attention and molecular events involved downstream of photoreceptor signaling remain to be identified.

In Arabidopsis, internode elongation reflects the activity of the basal part of the SAM, the rib zone. In light-grown plants, the rib zone is compact and mitotically inactive during vegetative growth, resulting in the formation of a compact rosette. At floral transition, the rib zone becomes activated to provide cells for rapid elongation of the internodes of the inflorescence stem (Vaughan, 1955; Sachs et al., 1959; Peterson and Yeung, 1972; Jacqmard et al., 2003; Bencivenga et al., 2016; Serrano-Mislata et al., 2017). Previously, ectopic expression of the Three-Amino Acid Loop Extension (TALE) homeobox gene *ARABIDOPSIS THALIANA HOMEOBOX GENE1* (*ATH1*) was shown to suppress growth of the inflorescence stem, due primarily to inhibition of internode elongation (Cole et al., 2006; Gómez-Mena and Sablowski, 2008; Rutjens et al., 2009; Ejaz et al., 2021). In wild-type plants, *ATH1* is expressed in the vegetative SAM, and its expression in this tissue is rapidly downregulated at floral transition, when stem growth is initiated. In addition, in plants lacking functional ATH1, the subapical region, where the rib zone is located, is enlarged during vegetative development, suggesting that ATH1 restricts growth of this part of the SAM (Proveniers et al., 2007; Gómez-Mena and Sablowski, 2008). In line with this, light-grown *ath1* mutants display slightly elongated rosette internodes, resembling those of higher-order photoreceptor mutants (Li et al., 2012; Ejaz et al., 2021). *ATH1* was originally identified in a screen for light-regulated transcription factor genes and its expression is induced by light during seedling de-etiolation (Quaedvlieg et al., 1995). In dark-grown seedlings of the photomorphogenic *cop1* mutant *ATH1* mRNA levels are elevated as well, suggesting that *ATH1* expression is under the control of COP1, a negative regulator of photomorphogenesis (Quaedvlieg et al., 1995; Proveniers et al., 2007). In line with this, *cop1* loss-of-function mutants exhibit a constitutive deetiolated phenotype in darkness, including formation of a compact rosette (Deng and Quail, 1992). Together with SUPPRESSOR OF PHYA-105 (SPA) proteins, COP1 forms an E3 ubiquitin ligase complex, which acts by regulating the stability of photomorphogenesis-promoting transcription factors. In addition, COP1/SPA stabilizes proteins of the PHYTOCHROME INTERACTING FACTOR (PIF) family in darkness to promote etiolation (Ponnu and Hoecker, 2021)). Upon exposure to light, phytochromes physically interact with PIF proteins and promote their turnover, resulting in de-etiolation (Pham et al., 2018a; Ponnu and Hoecker, 2021).

Here we show that ATH1 confers rosette habit in light-grown, vegetative Arabidopsis plants by integration of signals from multiple photoreceptors. In wildtype plants, *ATH1* expression can be induced by blue, red, and far-red light and this requires both the PHY and CRY families of photoreceptors. Dark-grown wildtype plants and higher-order photoreceptor mutants display strongly reduced levels of *ATH1* in the SAM and transgene expression of *ATH1* is sufficient to restore rosette habit in both cases. Finally, we introduce a regulatory feedback loop whereby multiple PIFs (i.e., PIF1, PIF3, PIF4, and PIF5) and ATH1 repress each other’s expression in a tissue-specific manner, contributing to the maintenance of rosette habit.

Furthermore, in the absence of light, *ATH1* expression can be induced by the direct availability of metabolic sugars to the SAM. We show that increasing amounts of sucrose result in a corresponding increase of *ATH1* expression and associated increased inhibition of vegetative internode elongation. Our data further show that both light- and metabolic signal-mediated induction of *ATH1* at the SAM requires activation of TOR kinase.

## Results

### Induction of *ATH1* expression restores a compact rosette habit in dark-grown seedlings

In higher plants, most of the above-ground part is generated by a combination of cell division, cell growth, cell expansion, morphogenesis, and differentiation at or near the SAM. Upon germination, the SAM becomes rapidly activated by light and starts a highly coordinated cell division program that continues throughout vegetative growth. In contrast, stem cells remain dormant when Arabidopsis plants are germinated and grown in darkness (Pfeiffer et al., 2016; Mohammed et al., 2017) As a result, dark-grown seedlings display a typical etiolated phenotype. This morphogenetic arrest can be overcome by the availability of sucrose to the aerial part of the plant. Sugar-induced dark morphogenesis of Arabidopsis plants follows the same developmental phases as light-grown plants. However, such plants fail to develop a compact rosette (Figure 1A, B) (Roldán et al., 1999; Mohammed et al., 2017).

**Figure 1:**
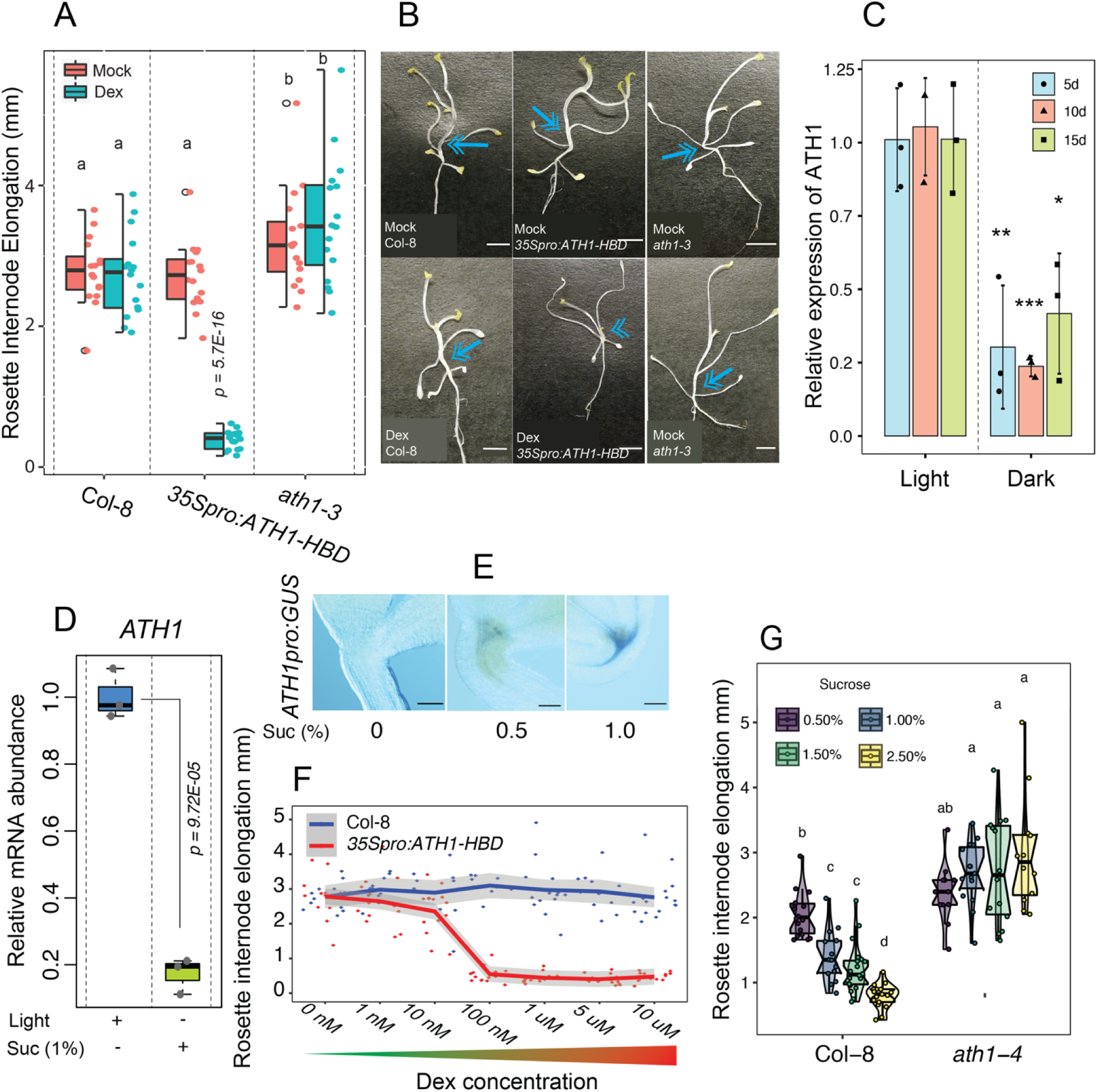
*ATH1* expression is sufficient to restore compact rosette growth in dark-grown Arabidopsis plants. **A:** Average rosette internode elongation in dark-grown Col-8, *ath1-3*, and *35Spro:ATH1-HBD* plants treated with 0.1% ethanol (mock) or 10 μM dexamethasone (Dex). Sucrose was added three days after the start of the experiment. **B:** Representative three-week-old, dark-grown plants used in A. Arrows indicate elongated rosette internodes, the arrowhead indicates suppression of internode elongation. Scale bars denote 2 mm. **C**: Relative expression of *ATH1* in Col-8 plants grown for either five, ten, or fifteen days (d) in continuous light or continuous darkness at 22°C. Sucrose was present at a one percent final concentration from the beginning of the experiment. Transcript levels were normalized to *MUSE3* (AT5G15400). The average of three biological replicates is shown. Error bars represent standard deviation of the ΔCT mean. P-values indicate significant differences; p-value < 0.05, p-value < 0.01, and p-value < 0.001 are represented by “*”, “**”, “***” respectively. **D**: Relative expression of *ATH1* in seven-day-old seedlings grown in continuous light (+) or continuous darkness (-) in the presence (+) or absence (-) of one percent sucrose. Transcript levels were normalized to *GAPC2* (AT1G13440). The average of three biological replicates is shown. The p-value for the observed difference is depicted in the figure. **E:** GUS-stained seven-day-old, dark-grown *ATH1pro:GUS* seedlings in the absence (0%) or presence (0.5% and 1%) of sucrose. GUS activity is visible as a blue precipitate. Scale bars represent 0.01 mm. **F**: Average rosette internode elongation in three-week-old dark-grown Col-8 and *35Spro:ATH1- HBD* plants treated with increasing concentrations of Dex (0 nM to 10 μM). **G**: Average rosette internode elongation of three-week-old dark-grown Col-8 and *ath1-4* seedlings treated with increasing concentrations of sucrose (0.5 to 2.5%). In **A** and **G**: Different letters indicate statistically significant differences (P< 0.05) as determined by one-way analysis of variance with Tukey’s honest significant difference post hoc test. In **A**: The p-value for the observed difference in *35Spro:ATH1-HBD* (mock and Dex) is depicted in the figure. In **A**, **F**, **G**: colored dots indicate rosette internode elongation scores of individual seedlings.

The compact rosette habit of light-grown Arabidopsis plants is conferred by ATH1 (Li et al., 2012; Ejaz et al., 2021). We tested whether *ATH1* expression is sufficient for development of a compact rosette in dark-grown plants. For this a fusion protein of ATH1 with the rat glucocorticoid receptor hormone-binding domain (HBD) under the control of the CaMV 35S promoter (*35Spro:ATH1- HBD*; Rutjens et al., 2009) was expressed in plants grown in continuous darkness in the presence of one percent sucrose (Figure 1A, B). Induction of nuclear expression of ATH1 using dexamethasone (Dex) resulted in strong repression of rosette internode elongation and, consequently, restoration of rosette habit in dark-grown plants, while Col-8 control plants and mock-treated *35Spro:ATH1-HBD* plants displayed elongated vegetative internodes, resulting in loss of rosette habit (Figure 1A, B). Surprisingly, in darkness in the presence of sucrose vegetative internodes of loss-of-function *ath1* mutants (Proveniers et al., 2007), were slightly more elongated than those in Col-8 wild-type control plants (Figure 1A, B), suggesting that *ATH1* might still be expressed to some extend in the absence of light, despite previous research indicating the absence of *ATH1* expression in dark-grown plants (Quaedvlieg et al., 1995).

Possibly, the addition of one percent sucrose to the growth medium, necessary to induce dark morphogenesis, results in induction of *ATH1* gene expression in the absence of light. To investigate whether *ATH1* is expressed in dark-grown seedlings in the presence of sucrose, we compared *ATH1* expression between light- and dark-grown seedlings, both supplied with one percent exogenous sucrose. As can be seen in Figure 1C, *ATH1* expression is significantly decreased but not entirely abolished in dark-grown seedlings. Compared to a light-grown situation, *ATH1* expression is more than 80% reduced (Figure 1D). Furthermore, in accordance with Quaedvlieg *et al*. (1995), *ATH1pro:GUS* analysis showed no *ATH1* induction in dark-grown plants in absence of sucrose, whereas *ATH1* promoter activity could be gradually induced by addition of increasing amounts of sucrose to the growth medium (Figure 1E). These findings indicate a dose-dependent relationship between sucrose concentration and ATH1 expression in dark-grown plants. Thus, in dark-grown Arabidopsis plants, sucrose can substitute for light to induce *ATH1* expression at the shoot apex.

Importantly, these observations suggest a close correlation between the amount of *ATH1* expressed at the shoot apex and the extent to which rosette internode elongation is suppressed. To test this, we analyzed elongation of vegetative internodes in dark-grown *35Spro:ATH1-HBD* plants exposed to increasing concentrations of Dex (Figure 1F). As Dex promotes nuclear accumulation of ATH1, increased Dex concentrations are expected to result in increased levels of nuclear ATH1 and, hence, stronger inhibition of internode elongation. This is exactly what was observed, with a maximum inhibitory effect on internode elongation in plants exposed to a Dex concentration of 100 nM, whereas none of the Dex concentrations tested had any effect on rosette internode elongation in control plants (Figure 1F). In line with these results, dark-grown Col-8 plants displayed increasing inhibition of rosette internode elongation when exposed to increasing concentrations of sucrose (0.5, 1.0, 1.5, or 2.5%). In these seedlings, rosette habit was completely restored in the presence of 2.5% sucrose, comparable to fully-induced dark-grown *35Spro:ATH1- HBD* plants (Figure 1A, F, G). As expected, in *ath1-4* mutant plants, internode elongation remained unaffected at all sucrose concentrations tested (Figure 1G). Taken together, this strongly suggests that sucrose-induced repression of rosette internode elongation in dark-grown plants is *ATH1*-dependent. These findings further show that typical loss of rosette habit, generally observed in sucrose-stimulated, dark-grown Arabidopsis plants, can be attributed to suboptimal *ATH1* expression at the shoot apex.

### SAM morphology of sucrose-stimulated, dark-grown seedlings strongly resembles that of light-grown *ath1* mutants

In light-grown *ath1* mutants elongation of vegetative internodes results from activation of stem development, as reflected by premature rib zone (RZ) activity (Roldán et al., 1999; Rutjens et al., 2009; Ejaz et al., 2021). To confirm that the elongated internode phenotype observed in dark-grown Arabidopsis plants also results from premature activation of stem development, we imaged shoot apices of both light- and dark-grown Col-8 seedlings and compared these with those of light-grown *ath1-4* seedlings (Figure 2A-C). We then measured the lengths of individual cells that make up a central cell file running from the L1 layer into the subapical RZ region of the SAM until where the hypocotyl vascular strands converge (Figure 2F). As expected, when grown for five days in continuous light, *ath1-4* mutants displayed elongated vegetative internodes, whereas those of Col-8 plants remained compact (Figure S2A, C). Comparing the *ath1-4* and Col-8 genotypes showed the first four cells of the central cell file to be of similar length. In contrast, more basal cells, in the RZ region, were significantly more elongated in *ath1-4* mutants (Figure 2A, C, F). This is in line with previous observations by Rutjens et al. (2009) who found that in *ath1* mutants RZ cells are more longitudinally elongated than in wild-type control plants. A similar SAM morphology was observed in Col-8 seedlings grown in darkness and supplied with one percent sucrose. Under these conditions, Col-8 rosette habit is no longer maintained, similar to light-grown *ath1* mutants, (Figure S2B). Compared to light-grown Col-8 seedlings, basal cells were significantly more elongated in dark-grown Col-8 seedlings in the presence of sucrose, whereas the lengths of the four apical central cells of light-grown Col-8 seedlings were not statistically different compared to dark-grown Col-8 (Figure 2A-C, F). Morphologically these basal cells resemble the elongated RZ cells of *ath1-4* mutants (Figure 2B, C).

**Figure 2:**
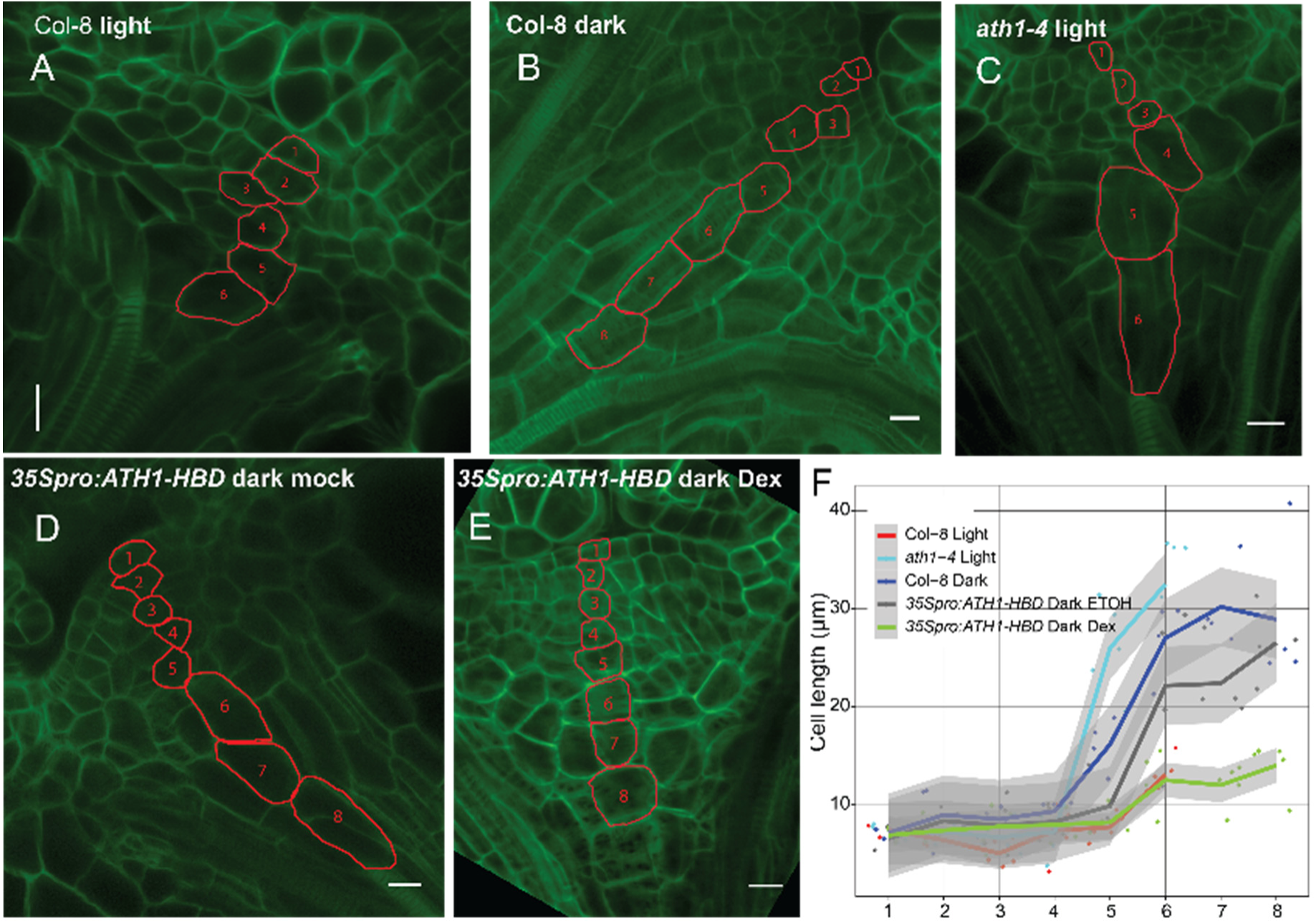
Sugar-induced dark-grown seedlings display a SAM morphology similar to light- grown *ath1* mutants. Median longitudinal optical sections through the shoot apical meristems of five-day-old Col-8 (**A, B**), *ath1-4* (**C**), and *35Spro:ATH1-HBD* (**D, E)** seedlings grown at 27°C in the presence (**A**, **C**) or absence (**B**, **D**, **E**) of light. Mock treatment (**D**) is 0.1% ethanol, Dex treatment (**E**) is 10 μM dexamethasone. Cells marked in red form a central cell file extending from the epidermis into the subapical region that forms the rib zone. Scale bars represent 10 µm. **F**: Quantification of cell lengths as illustrated in **A**-**E**. Individual cell lengths were measured per position in apical-basal direction. Per genotype and condition four or five individual apices were analyzed. The numbers on the x-axis correspond to the cell position as depicted in **A-E**.

Since ectopic expression of *ATH1* restored a compact rosette habit in dark-grown seedlings (Figure 1A, B; S2D, E), we examined whether this is caused by inhibition of RZ activity. While dark-grown mock-treated *35Spro:ATH1-HBD* plants showed a SAM morphology similar to that of dark-grown Col-8 control plants, induction of *ATH1* specifically repressed cell elongation in the basal RZ cells (Figure 2B, D-F). Taken together, these findings indicate that loss of rosette habit in Arabidopsis plants in the absence of light, results from premature RZ activation due to significantly reduced *ATH1* expression at the shoot apex. Moreover, we noted that in dark-grown seedlings the central cell file always contained more cells than that of light grown plants (Figure 2A-F). This seems to be an effect of light independent of ATH1 as this was found for all genotypes analyzed.

### ATH1 functions downstream of multiple photoreceptors to maintain a compact rosette habit

Loss of rosette habit due to elongation of vegetative internodes, reminiscent of the caulescent growth habit of sucrose-induced, dark-grown Arabidopsis plants, can also be observed in light-grown photoreceptor mutants. Elongation of vegetative internodes is clearly visible in higher order mutants, such as *phyA phyB* (*phyAB*), *phyA phyB phyD* (*phyABD*), *phyA phyB phyE* (*phyABE*), *phyB phyD phyE* (*phyBDE*), *phyA phyB phyD phyE* (*phyABDE*) mutants, a quintuple *phy* mutant, and *phyB cry1* mutants (Devlin et al., 1996; Devlin et al., 1998; Whitelam et al., 1998; Devlin et al., 1999; Mazzella et al., 2000; Devlin et al., 2003; Strasser et al., 2010; Li et al., 2012). *ATH1* expression is strongly light-dependent (Quaedvlieg et al., 1995; Roldán et al., 1999; Proveniers et al., 2007) (Figure 1C, D), raising the question of whether light-mediated expression of *ATH1* depends on these photoreceptors. If so, it is of interest to investigate whether reduced levels of *ATH1* can explain the significant elongation of internodes between adjacent rosette leaves displayed by the respective photoreceptor mutants. To answer these questions, we first grew wild-type seedlings under different wavelengths of light and analyzed *ATH1* mRNA levels (Figure 3A). This revealed that apart from white light, as reported before, monochromatic blue, red, and far-red light induce *ATH1* expression to significant levels, suggesting that *ATH1* is under control of multiple photoreceptors.

**Figure 3:**
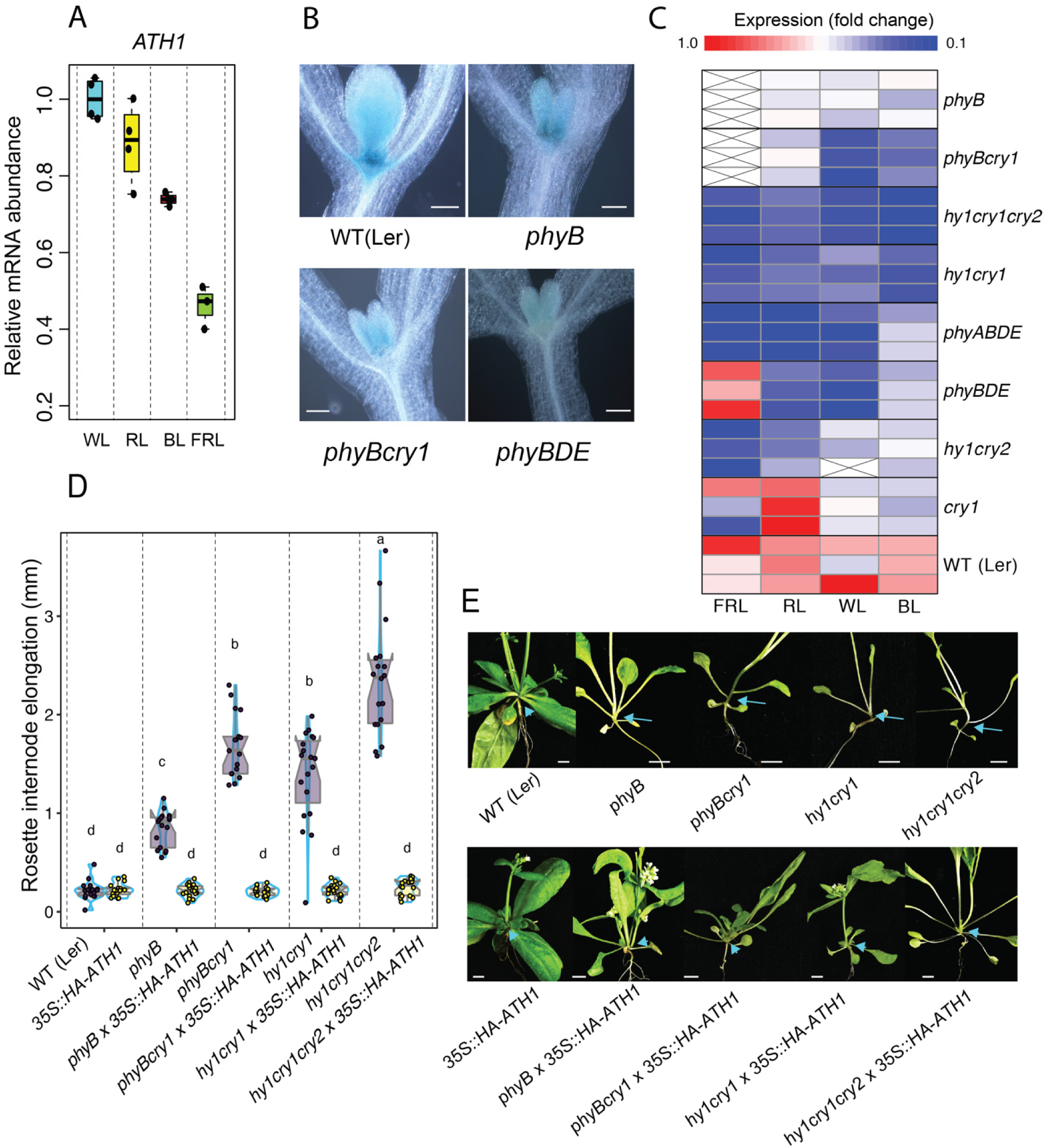
Significant reduction in *ATH1* expression levels underlies loss of rosette habit in photoreceptor mutants. **A**: Expression of *ATH1* in seven-day-old seedlings (L*er*) grown in SD white light (WL), red (RL), blue (BL), far-red light (FRL) or continuous darkness (DRK). Transcript levels were normalized to *GAPC2* (AT1G13440). Dots indicate the average values of four biological replicates per light treatment, each consisting of 40-50 seedlings. **B**: Shoot apices of GUS-stained, seven-day-old ATH1_pro_:GUS seedlings in different genetic backgrounds (Col-8, *phyB*, *phyBcry1*, and *phyBDE*). Plants were grown in whit light under short- day conditions. Scale bars represent 0.01 mm. **C**: Heat map generated from qPCR data on relative *ATH1* expression in indicated photoreceptor mutants (see **Figure S1**), when compared to wild-type control plants (L*er*). Transcript levels were normalized to *GAPC2* (AT1G13440; BL) or *MUSE3* (AT5G15400; RL, FRL and WL). The average of three biological replicates is shown. At least 20 shoot apices were used for each biological replicate. Red corresponds to high relative expression and dark blue corresponds to low relative expression. A linear fold change scale is displayed on top. **D:** Average rosette internode elongation in WT (L*er*), *phyB*, *phyBcry1*, *hy1cry1*, and *hy1cry1cry2* in the absence or presence of a *Pro_35S_:HA-ATH1* transgene. Plants were grown under LD conditions. Different letters denote statistically significant differences between groups (P < 0.05) as determined by 1-way ANOVA followed by Tukey’s post hoc test. Colored dots indicate the average rosette internode length per individual (n ≥16 individual plants per genotype). **E:** Representative plants from **D**. Arrows indicate elongated rosette internodes, the arrowheads indicate complete suppression of internode elongation. Scale bars represent 5 mm.

We next determined *ATH1* promoter activity and mRNA levels in a series of phytochrome and/or cryptochrome photoreceptor mutants grown under various light quality conditions (Figure 3C; S1). Whereas *ATH1* transcript levels in response to white light were somewhat decreased in *phyB* and *cry1* single mutants, combination of both mutations significantly affected white light-mediated induction of *ATH1* expression (Figure 3C; S1). Similarly, introduction of additional *phy* mutations in a *phyB* background or the combination of the phytochrome chromophore biosynthesis mutant *hy1* with *cry1* and/or *cry2* mutations resulted in moderate to severe reduction in *ATH1* transcript levels in white light, confirming that light-mediated regulation of *ATH1* is under control of multiple photoreceptors (Figure 3C, S1A). Repeating experiments under monochromatic red, far-red, or blue-light conditions revealed that red-light-mediated induction of *ATH1* expression is mostly the result of phyB function, in cooperation with phyD and phyE (Figure 3C, S1B), whereas phyA is largely responsible for *ATH1* induction in far-red light (Figure 3C, S1D). Under monochromatic blue light, CRY1 and CRY2 redundantly contribute to *ATH1* gene activity, with CRY1 being the predominant cryptochrome under the conditions tested (Figure 3C, S1C). We further noticed that all photoreceptor mutants previously reported to have a loss of rosette habit, including *phyBDE* and *phyB cry1* mutants (Mazzella et al., 2000; Li et al., 2012), all had severely reduced levels of *ATH1* (Figure 3C). Internode elongation reflects activity of the basal part of the SAM, the rib zone, and ATH1 controls activity of this tissue. We compared the spatial activity of the *ATH1* promoter in these higher order photoreceptor mutants with that in L*er* control plants and a *phyB* mutant. In line with previous observations (Proveniers et al., 2007), high levels of GUS activity were present in the SAM and emerging leaf primordia of L*er ATH1pro:GUS* seedlings grown in white light. Corroborating our qPCR data, GUS activity was significantly reduced in *phyB cry1 ATH1pro:GUS* and *phyBDE ATH1pro:GUS* plants, whereas in a *phyB* background GUS activity was only moderately affected (Figure 3B). Interestingly, the largest effect of reduced photoreceptor signaling on *ATH1* promoter activity was in the SAM region. In both *phyB cry1* and *phyBDE* backgrounds hardly any residual GUS activity could be detected in the SAM, including the rib zone area, whereas in leaf primordia a more modest reduction in *GUS* expression was observed (Figure 3B). Taken together, these findings suggest that phytochrome and cryptochrome photoreceptor families contribute to rosette growth habit in light-grown vegetative Arabidopsis plants through induction of *ATH1* expression in the SAM.

To further test this hypothesis, we constitutively expressed *ATH1* (*Pro35S:HA-ATH1*) in a number of photoreceptor mutants that display elongation of vegetative internodes when grown under standard, long-day conditions, with partial to complete loss of rosette growth habit as a result (Figure 3D, E). Under these growth conditions, L*er* control plants never display any detectable elongation of rosette internodes. In contrast, internodes of *phyB*, *phyBcry1*, *hy1cry1*, and *hy1cry1cry2* mutants were visibly elongated (Figure 3D, E). Strikingly, the extent to which rosette internode elongation was affected in these genotypes nicely correlates with earlier observed *ATH1* expression levels in the respective mutant backgrounds (Figure 3C-E; S1A). As expected, constitutive expression of *ATH1* completely suppressed elongation of vegetative internodes in all four mutants analyzed (Figure 3D, E). Thus, reestablishing high levels of *ATH1* expression is sufficient to restore a compact rosette habit in higher order photoreceptor mutants.

In conclusion, rosette growth habit, quintessential for light-grown vegetative Arabidopsis plants, is imposed by *ATH1* activity in the shoot apex under control of multiple blue and red/far-red light photoreceptors.

### Light-mediated regulation of *ATH1* expression is controlled by central light signaling components

*ATH1* was first identified in our lab as a light-regulated transcription factor gene that is derepressed in dark-grown *cop1* mutants (Quaedvlieg et al., 1995). COP1, in conjunction with SPA proteins, functions as a repressor of light signaling in darkness. In the light, phytochrome family members, activated by red or far-red light (in case of phyA) wavelengths, and cryptochromes 1 and 2, activated by blue light wavelengths, suppress the activity of the COP1/SPA complex to promote photomorphogenesis (Ponnu and Hoecker, 2021). Light-mediated *ATH1* expression can be induced by all three light qualities, involving both phytochrome and cryptochrome family members. Since COP1 is a downstream signaling component of these photoreceptor families, we analyzed the role of COP1 in the regulation of *ATH1* expression and rosette growth habit. First, we compared *ATH1* expression levels in dark-grown Col-8 seedlings to those in dark-grown *cop1- 4* mutants, both in the presence and absence of sucrose (Figure 4A). As shown before, in Col-8 *ATH1* expression can be induced in darkness by the addition of sucrose to the growth medium to a final concentration of one percent, whereas in the absence of sucrose transcript levels remain low. In line with the observations by Quaedvlieg et al. (1995), in dark-grown *cop1-4* mutants, carrying a mild loss-of-function allele of *COP1*, *ATH1* expression was clearly derepressed. Already in the absence of sucrose, *ATH1* mRNA levels accumulated to levels higher than those observed in Col-8 control plants supplied with one percent sucrose. In the presence of sucrose, *cop1-4 ATH1* transcript levels increased even further (Figure 4A), indicating that light and sucrose signaling contribute, at least partially, independently to induce of *ATH1* gene expression.

**Figure 4:**
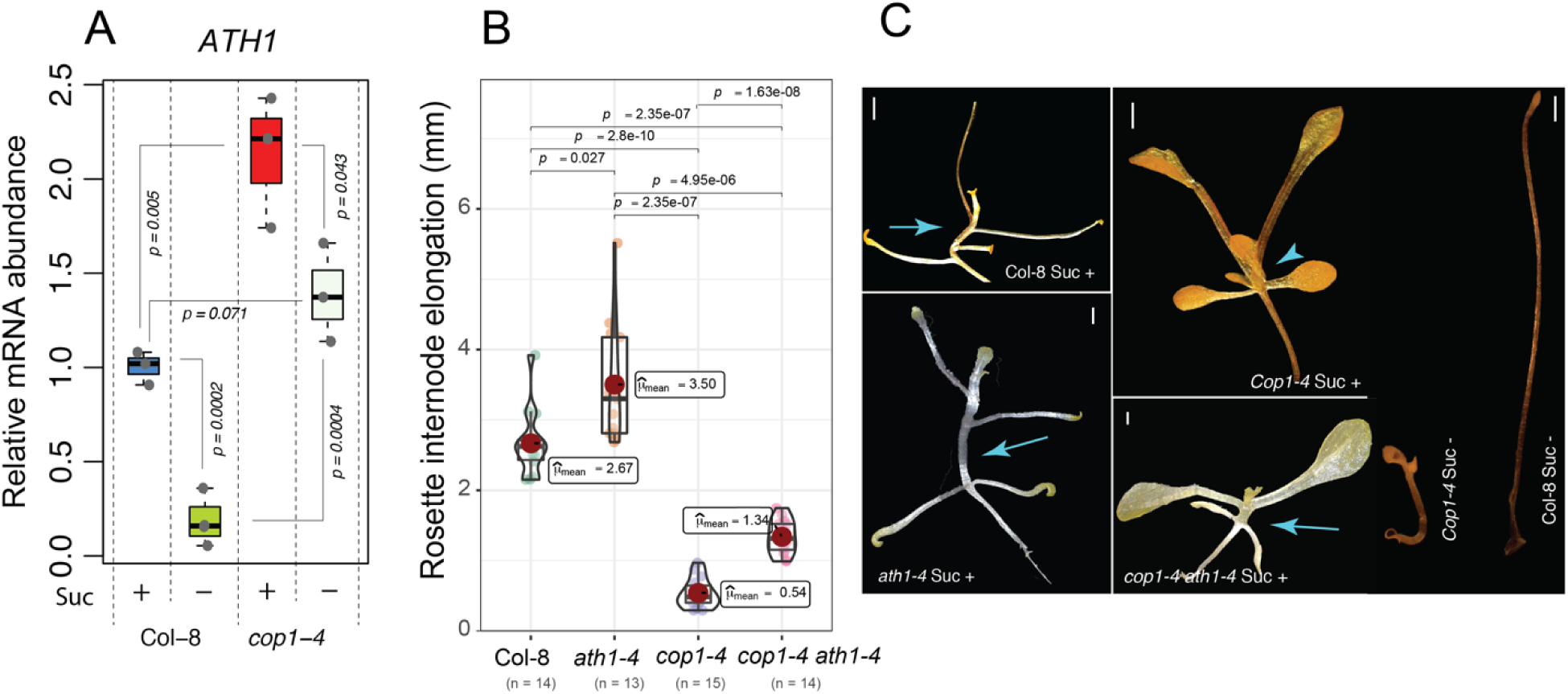
Derepression of *ATH1* contributes to a compact rosette habit in dark-grown *cop1* mutants. **A**: Relative mRNA abundance of *ATH1* in shoot apices of two-week-old dark-grown seedlings. The average of three biological replicates is shown. At least 20 shoot apices were used for each biological replicate. The p-value of significant differences, as determined by the two-tailed Student’s t-test, are indicated in the figure. **B:** Average rosette internode lengths of 3-week-old Col-8, *ath1-4*, *cop1-4,* and *cop-1 ath1-4* plants (n ≥13) grown in continuous darkness for three weeks at 22°C, in the presence of one percent sucrose. The significant differences are indicated in the figure as determined by Pairwise test Games- Howell according to R package ggstatsplot (Patil, 2021). **C**: Representative plants from **B**. Arrows indicate elongated rosette internodes, arrowheads indicate complete suppression of internode elongation. Scale bars represent 0.5 mm. Sucrose (Suc +) or sorbitol (Suc -), both to a final concentration of one percent, were added three days after start of the experiment (**A**, **C**)

Consistent with these observations, vegetative internodes of dark-grown *cop1-4* mutants supplemented with one percent sucrose failed to elongate, contrasting to those of Col-8 control plants. As a result, *cop1-4* plants adopted a compact rosette growth habit, reminiscent of that seen in dark-grown, induced *35Spro:ATH1-HBD* plants supplemented with sucrose (Figure 4B, C; 1B). The compact rosette habit in darkness is lost in *cop1-4 ath1-4* double mutants, indicating that this photomorphogenesis phenotype requires ATH1 activity (Figure 4C). However, when compared to *ath1-4* single mutants, vegetative internode elongation in *cop1 ath1* double mutants is much reduced, suggesting the involvement of other loci apart from *ATH1* (Figure 4B, C). In the absence of sucrose, both Col-8 and *cop1-4* seedlings failed to develop beyond the seedling stage in the dark (Figure 4C), most likely due to the lack of stem cell activation at the shoot apex (Pfeiffer et al., 2016).

In addition to COP1, PIFs also function downstream of both phytochrome and cryptochrome light signaling (Ma et al., 2016; Pedmale et al., 2016; Pham et al., 2018b). PIF-family transcription factor proteins function as negative regulators of light responses, in a partially redundant manner, to maintain skotomorphogenesis in dark-grown seedlings. Upon exposure to light, phytochromes promote the turnover of PIFs, whereas photoactivated CRY1 has been shown to interact with PIF4, resulting in suppression of the transcriptional activity of PIF4 (Ma et al., 2016). As a consequence, plants switch from skotomorphogenesis to photomorphogenesis. In line with this, the quadruple *pif1pif3pif4pif5* (*pifq*) mutant displays a constitutively photomorphogenic phenotype in darkness (Leivar et al., 2009). In addition to their role as transcription factor proteins, PIFs can directly interact with COP1, thereby enhancing substrate recognition and ubiquitination activity of the COP1 E3 ligase complex (Xu et al., 2014; Kathare et al., 2020). Therefore, we tested whether PIF family members act as upstream mediators of *ATH1* gene expression in the regulation of rosette habit in Arabidopsis. To this end, we first analyzed rosette habit in a series of dark-grown single, double, triple, and quadruple *pif* mutant combinations in the presence of one percent sucrose. Under these conditions, Col-8 control plants display a clearly visible elongation of vegetative internodes, resulting in loss of rosette habit. In contrast, in the quadruple *pifq* mutant complete repression of rosette internode elongation was observed, resulting in the formation of a compact rosette in the absence of light (Figure 5A). Of the single *pif* mutants tested, only *pif4* displayed a significant reduction in rosette internode length (37% less elongation) when compared to control plants, whereas *pif3* and *pif7* mutants were largely unaffected. None of the double (*pif1pif3*, *pif3pif4*, *pif4pif5*) or triple mutants (*pif1pif3pif4*, *pif1pif3pif5*, *pif3pif4pif5*) were as compact as the *pifq* mutant (Figure 5A), indicating that PIF1, PIF3, PIF4, and PIF5 redundantly contribute to elongation of rosette internodes in etiolated plants. Of these four proteins, PIF4 contributes the most, as can be inferred from its single mutant phenotype and the significant inhibition of internode elongation in all higher order mutants carrying a *pif4* allele, while inhibition of internode elongation is absent in *pif1pif3* mutants and only subtly enhanced by *pif5* in *pif4pif5* and *pif3pif4pif5* mutants (Figure 5A).

**Figure 5:**
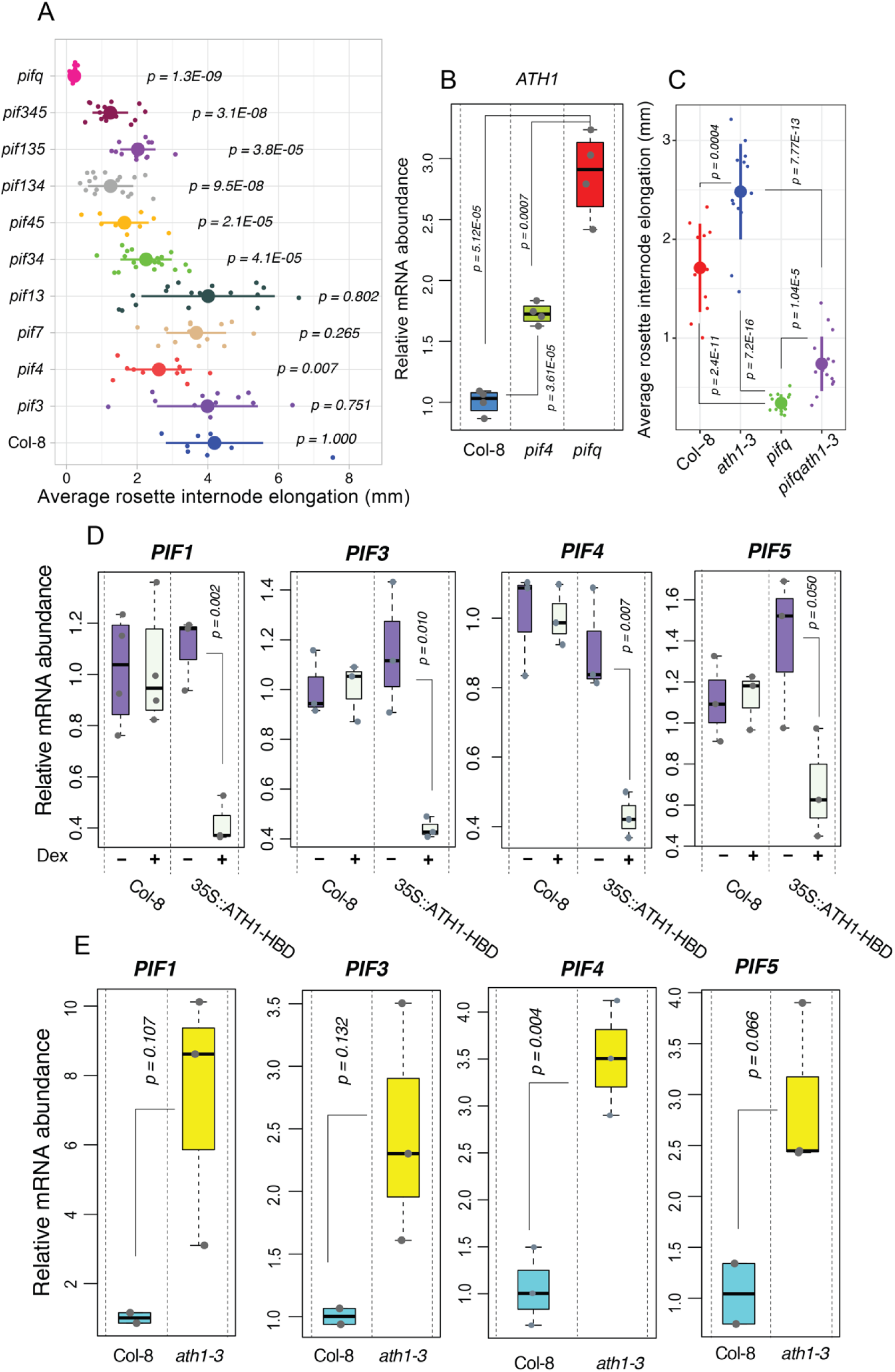
A double-negative feedback loop between ATH1 and PIFs is required for initiation and maintenance of rosette growth habit. **A**: Average internode lengths of 3-week-old Col-8, *pif3*, *pif4*, *pif7*, *pif1pif3*, *pif3pif4*, *pif4pif5*, *pif1pif3pif4*, *pif1pif3pif5*, *pif3pif4pif5*, and *pifq* (*pif1pif3pif4pif5*) plants grown in continuous darkness at 22°C. Sucrose was added to the medium to a final concentration of one percent three days after the start of the experiment. Colored dots indicate average rosette internode elongation scores of individual seedlings (n ≥9). *The p-value* of significant differences compared to wild type Col-8, as determined by the two-tailed Student’s t-test, are indicated in the figure. **B**: Relative mRNA abundance of *ATH1* in SAM-enriched tissue of 14-day-old, dark-grown Col- 8, *pif4*, and *pifq* seedlings (n≥20 per biological replicate; four biological replicates). Transcript levels were normalized to *GAPC2* (AT1G13440). The p-value of significant differences, as determined by the two-tailed Student’s t-test, are indicated in the figure. Sucrose was present at a one percent final concentration from the start of the experiment. **C**: Average internode lengths of three-week-old Col-8, *pif4*, *pifq*, and *pifq ath1-3* plants grown in continuous darkness at 22°C. Sucrose was added to the medium to a final concentration of one percent three days after the start of the experiment. The p-value of significant differences, as determined by the two-tailed Student’s t-test, are indicated in the figure. Colored dots indicate average rosette internode elongation scores of individual seedlings (n ≥11). **D**, **E**: Relative mRNA abundance of indicated *PIF* genes in SAM-enriched tissue of 14-day-old, dark-grown Col-8 and *35Spro:ATH1-HBD* (n ≥3) (**D**) or 39-day-old, light-grown (SD conditions) Col-8 and *ath1-3* seedlings (n ≥2) (**E**). For (**D**) seedlings were treated with a mock (0.1% ethanol, Dex -) or 10 μM dexamethasone (Dex +) at day three, and in total, 30-40 shoot apices were used for each biological replicate (**D**). For light-grown plants (**E**), three shoot apices were used per biological replicate. Transcript levels were normalized to *GAPC2* (AT1G13440). Seedlings were treated with a mock (0.1% ethanol, Dex -) or 10 μM dexamethasone (Dex +) (**D**). The p-value of significant differences, as determined by the two-tailed Student’s t-test, are indicated in the figures.

The prominent role of PIFs in the suppression of photomorphogenesis and the striking resemblance of *pifq* mutants to DEX-induced *35Spro:ATH1-HBD* plants when grown in darkness (Figure 1A, B), prompted us to investigate whether PIF proteins are upstream regulators of *ATH1*. ATH1 functions in the shoot apex to control internode elongation. Therefore, *ATH1* transcript levels were compared between shoot apices of 14-day-old, dark-grown Col-8, *pif4* and *pifq* plants (Figure 5B). A significant increase in *ATH1* mRNA levels was observed in both *pif4* (1.7x higher) and *pifq* (3x higher) mutants when compared to control plants, in line with the observed effects on inhibition of internode elongation. To examine whether this increase in *ATH1* expression is responsible for the formation of the compact rosette habit observed in *pifq* mutants when grown in darkness, we introduced the *ath1-3* loss-of-function allele in the *pifq* mutant background and tested the resulting plants for rosette internode elongation when grown in darkness in the presence of one percent sucrose (Figure 5C). Under these conditions *pifq* mutants grow a compact rosette, whereas rosette habit is lost in Col-8 control plants and *ath1* mutants, with the latter having the most elongated internodes (Figure 5A, C). Surprisingly, vegetative internodes of *pifq ath1* plants were only mildly elongated, resulting in a partial loss of rosette habit. Compared to *ath1* plants, internodes of *pifq ath1* were on average 70% shorter (Figure 5C). This unexpected result suggests that PIFs control rosette internode elongation mostly independent of ATH1 or that the relationship between PIFs and ATH1 is more complex. Recently, PIF4 has been identified as a direct binding target of ATH1 (Ejaz et al., 2021). In this study no significant differences in *PIF4* expression could be detected between *ath1* and WT plants when analyzed on a whole-seedling basis. However, this does not rule out the presence of a regulatory feedback loop between PIFs and ATH1 at a tissue-specific level. Such ATH1-PIFs feedback regulation could explain the observed internode phenotype in *pifq ath1* mutants. To explore the presence of such regulatory interaction between ATH1 and PIFs, we quantified expression of PIF family genes in genotypes with altered expression of *ATH1*. We specifically focused on PIF gene expression in shoot apices (Figure 5D, E) as ATH1 activity in the shoot apex underlies its inhibitory effect on internode elongation and PIFs are ubiquitously expressed (Wu et al., 2020) In sucrose-supplied, dark-grown plants *ATH1* expression levels are low (Figure 1C, D) and ATH1 is expected to have an inhibitory effect on *PIF* expression. Therefore, inducible *35Spro:ATH1-HBD* plants were used to examine the effect of ATH1 on *PIF1*, *PIF3*, *PIF4* and *PIF5* mRNA levels in dark conditions (Figure 5D). In light-grown vegetative plants, *ATH1* expression levels at the shoot apex are relatively high. Therefore, in light conditions the effect of ATH1 on gene expression of these four *PIFs* was analyzed using *ath1-3* plants (Figure 5E). In both conditions, a clear effect of ATH1 on *PIF* gene expression was observed. In dark-grown plants, induction of ATH1 accumulation in the nucleus resulted in significant down- regulation of *PIF1*, *PIF3* and *PIF4*, and, to a lesser extent, *PIF5* expression (Figure 5D). On the other hand, in light-grown plants mRNA levels of *PIF1*, *PIF3*, *PIF4*, and *PIF5* were up-regulated in the absence of *ATH1*. In *ath1* plants, PIF1 levels in the shoot apex were up to 9 times higher when compared to control plants, whereas *PIF3*, *PIF4*, and *PIF5* levels were 2.5-3 times increased (Figure 5E). When analyzed on a whole-plant level, under both conditions no significant differences in *PIF* expression levels could be detected between either *35Spro:ATH1-HBD* or *ath1- 3* plants and control plants (data not shown). Together, these results indicate that ATH1 acts as a negative regulator of, at least, *PIF1*, *PIF3*, *PIF4*, and *PIF5* gene expression, in a tissue-specific manner. These data support the presence of a regulatory feedback loop at the transcriptional level between ATH1 and PIF family members where ATH1 and PIFs repress each other’s expression. This ATH1-PIF interdependence for suppression of rosette internode elongation could explain the partial loss of rosette habit in *cop1-4 ath1-4* double mutants observed earlier (Figure 4B, C), since functional PIF1, PIF3, PIF4 and PIF5 proteins are required for dark-mediated rosette internode elongation in the absence of ATH1 (Figure 5C) and these PIF proteins are degraded in darkness in the presence of a *cop1-4* mutation (Pham et al., 2018c; Pham et al., 2018a).

Overall our data show that loss of rosette habit in dark-grown Arabidopsis plants is part of a skotomorphogenic developmental program, achieved through active repression of *ATH1* gene expression, mediated by COP1 and PIF family members.

### Photosynthetic sugars are not a prerequisite for light-induced *ATH1* expression

*ATH1* expression in the shoot apex can be induced both by light and sucrose (Figure 1D, E; 3A). Since light not only acts as a developmental signal, but also as an energy source through photosynthesis, we wondered about the exact role of light in the induction of *ATH1* gene expression. Therefore, we examined *ATH1* promoter activity in plants where photosynthesis was chemically inhibited. To this end, *ATH1pro:GUS* seedlings were grown in darkness for five days, without metabolizable sugar in the medium to deplete plant metabolizable sugar. Five hours before light treatment, either norflurazon or lincomycin repressors of photosynthesis were added to the medium, after which plants were grown for an extra two days in continuous light (Figure 6A). Compared to mock treatment (0.1% ethanol), inhibition of photosynthesis by chemical interference resulted in slightly reduced *ATH1* expression (Figure 6B). To avoid potential indirect effects of these pharmacological treatments, we also examined the impact of CO_2_ withdrawal on *ATH1* expression. Removal of CO_2_ from the atmosphere, through chemical absorption by NaOH + CaO, inhibits photosynthetic carbon assimilation and, thereby, the accumulation of metabolizable sugars. Similar to the pharmacological treatments in the absence of photosynthesis, *ATH1* promoter activity was decreased, but GUS staining was still clearly visible (Figure 6B; S3A). This indicates that *ATH1* gene expression is affected by light acting both as a developmental trigger and as a source of metabolizable sugars through photosynthesis. It further shows that sucrose produced through photosynthesis contributes to, but is not a prerequisite for light-induced *ATH1* expression. This is in line with the observation that *ATH1* expression is derepressed in dark-grown *cop1* seedlings even in the absence of sucrose (Figure 4A).

**Figure 6:**
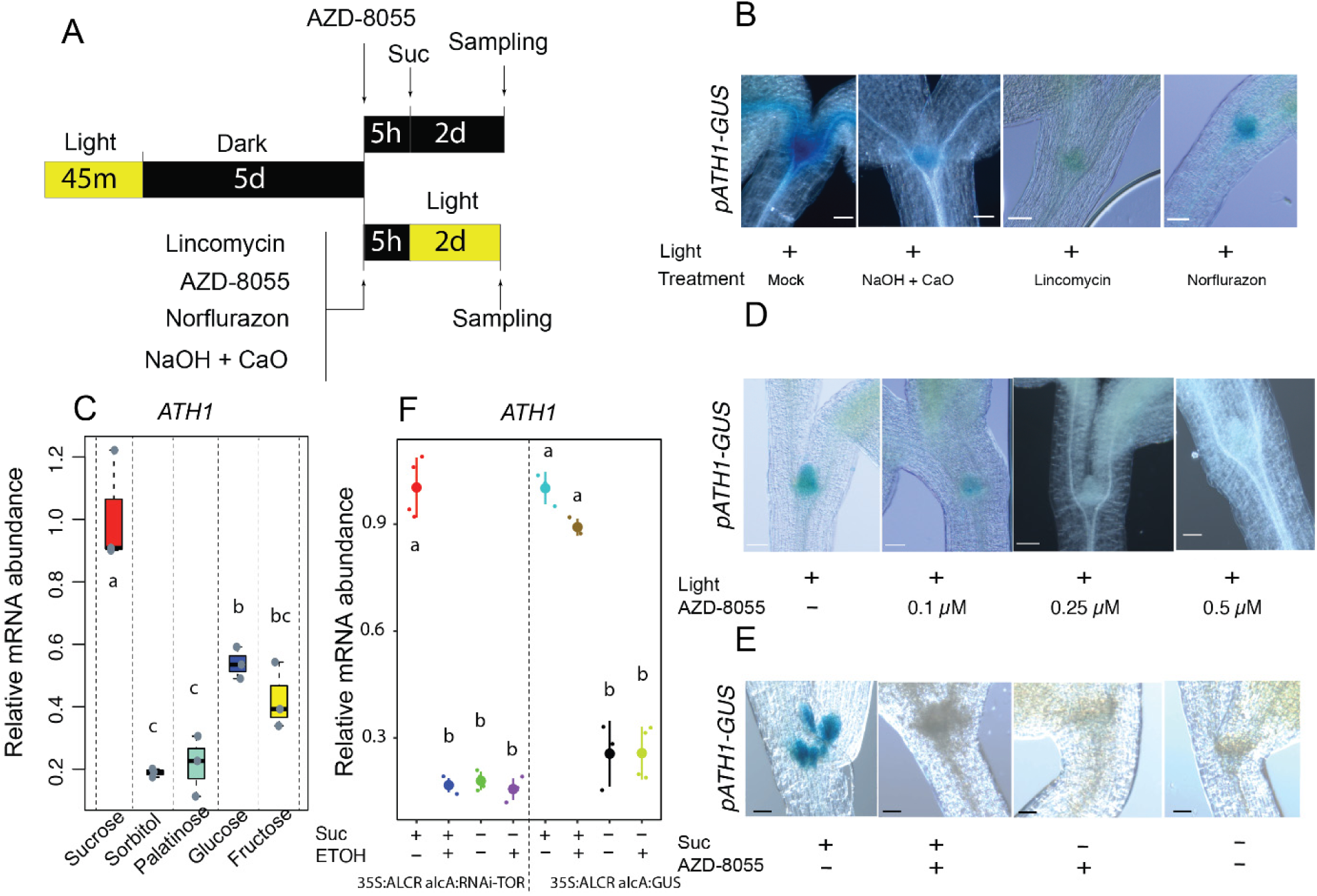
*ATH1* expression is independently regulated by light and sucrose. **A**: Schematic representation of the experimental setup. *ATH1_pro_:GUS* seeds were light-treated for 45 minutes to stimulate germination, before growth in continuous darkness for five days. AZD- 8055, lincomycin, norflurazon, or NaOH + CaO were then added and seedings were grown for an additional five hours in darkness before switching to *ATH1*-inducing conditions (continuous light or continued growth in darkness in the presence of sucrose (Suc)) for two more days. **B**: Shoot apices of GUS-stained *ATH1_pro_:GUS* seedlings treated with either NaOH + CaO, 0.5 mM lincomycin, or 5 µM norflurazon according to the scheme depicted in (**A**). Mock-treatment was with ethanol to a final concentration of 0.1%. Scale bars represent 0.05 mm. **C**: Relative expression of *ATH1* in seven-day-old, dark-grown seedlings, grown in the presence of either sucrose, glucose, fructose, palatinose, or sorbitol, all at a final concentration of one percent in the growth medium. Sugars were added at the start of the experiment. Transcript levels were normalized to *MUSE3* (At5g15400). The average of three biological replicates is shown. At least 30 seedlings were used for each biological replicate. **D, E**: Shoot apices of GUS-stained *ATH1_pro_:GUS* seedlings treated with the TOR kinase inhibitor AZD-8055 before switching to ATH1-inducing conditions (continuous light (**D**) or darkness in the presence of one percent sucrose (**E**)) according to the scheme depicted in (**A**). Scale bars represent 0.01 mm. **F**: Relative expression of *ATH1* in seven-day-old, dark-grown *35S::ALCR alcA:RNAi-TOR* and *35S:ALCR alcA:GUS* (control line) seedlings in the presence or absence of ethanol (ETOH; 0.1%) and/or sucrose (Suc; 1%). Experimental setup was as in (**A)**. ETOH was added after five days of growth in darkness. After an additional five hours in darkness sucrose was added and plants were sampled after two more days in darkness. Transcript levels were normalized to *GAPC2* (AT1G13440). The average of three biological replicates is shown. At least 30 seedlings were used for each biological replicate. Different letters denote statistically significant differences between groups (P < 0.05) as determined by 1-way ANOVA followed by Tukey’s post hoc test.

Sugars function as energy resource and as signaling molecules (Li and Sheen, 2016). To distinguish between these two functions in the induction of *ATH1* expression at the SAM Col-8 plants were grown in darkness in the presence of either sorbitol, sucrose, glucose, fructose, or palatinose, followed by quantification of *ATH1* mRNA levels and promoter activity (Figure 6C; S3C). Sorbitol and palatinose are both non-metabolizable sugars. Palatinose is a metabolically inactive structural isomer of sucrose that was shown to function as a signaling molecule, whereas sorbitol is neither metabolized nor a signaling molecule (Ramon et al., 2008). Glucose, fructose and sucrose are metabolizable sugars also known to function as signaling molecules (Rabot et al., 2012). Neither sorbitol nor palatinose addition resulted in a significant induction of *ATH1* expression, whereas a clear increase in *ATH1* could be observed when either sucrose, glucose or, to a lower extent, fructose was present in the growth medium (Figure 6C). These findings strongly suggest that sugars as energy source induce *ATH1* expression.

### Sucrose and light independently regulate *ATH1* expression via TOR kinase

TOR kinase, a critical sensor of resource availability, is required for the activation of both shoot and root apical meristems (Xiong et al., 2013; Pfeiffer et al., 2016; Li et al., 2017). TOR kinase integrates, among others, energy and environmental cues, including light signals to direct growth and development. The fundamental role of TOR kinase in coordinating growth and development downstream of light and energy signals, led us to investigate whether induction of *ATH1* expression involves TOR kinase activity. Employing a similar experimental setup as mentioned in the previous section, the effect of the ATP-competitive TOR kinase inhibitor AZD-8055 on light- and sucrose-induced *ATH1* promoter activity was studied. AZD-8055 was added to 5-day-old, dark-grown *ATH1pro:GUS* seedlings. After 5 hours either sucrose was added to the growth medium and plants were left to continue growing in darkness, or plants were shifted to continuous light conditions for 48 hours (Figure 6A). Light-mediated induction of *ATH1* promoter activity was efficiently suppressed by AZD-8055, resulting in complete inhibition of promoter activity at a concentration 0.5 µM (Figure 6D). Similarly, addition of AZD-8055 fully inhibited the positive effect of sucrose on ATH1 promoter activity at the shoot apex (Figure 6E; S3B).

AZD-8055 is a selective and potent TOR inhibitor (Montané and Menand, 2013; Dong et al., 2015), but the use of pharmacological agents poses a risk of non-anticipated side effects. Loss-of-function *tor* mutants are embryo lethal (Menand et al., 2002), and therefore the previously reported ethanol-inducible TOR-RNAi line, *35S:ALCR alcA:RNAi-TOR* and its control line *35S:ALCR alcA:GUS* (Deprost et al., 2007), were used to further study the effects of reduced TOR kinase activity on *ATH1* gene expression. In line with our AZD-8055 experiments, conditional silencing of the AtTOR gene by ethanol in *35S:ALCR alcA:RNAi-TOR* seedlings led to complete inhibition of sucrose-mediated induction of *ATH1* expression (Figure 6F). These results imply that light and metabolic signals converge on TOR kinase in controlling induction of *ATH1* expression at the SAM.

Recently, it was reported that TOR kinase and downstream signaling contribute to the de-etiolation process and that COP1 represses TOR activity during skotomorphogenic development (Chen et al., 2018). We therefore tested whether derepression of *ATH1* expression in dark-grown *cop1* mutants is also TOR dependent. QPCR analysis revealed that this is indeed the case, as in the presence of AZD-8055 *ATH1* expression is no longer derepressed in dark-grown *cop1-4* plants (Figure S4).

In conclusion, these findings suggest TOR kinase integrates light and sucrose signals leading to activation of *ATH1* gene expression at the shoot apex (Figure 7). Upstream of TOR kinase, a PHY- COP1 regulatory pathway functions as a negative regulator of TOR activity. When grown in darkness, COP1 inhibits TOR, resulting in repression of *ATH1*. As a consequence, in dark-grown plants compact rosette habit is lost due to activation of stem development, resulting in the elongation of vegetative internodes. In the light, COP1 activity is inhibited allowing for TOR kinase to induce *ATH1* expression in the shoot apex as part of the deetiolation process, giving rise to the compact rosette habit that is characteristic for *A. thaliana*.

**Figure 7:**
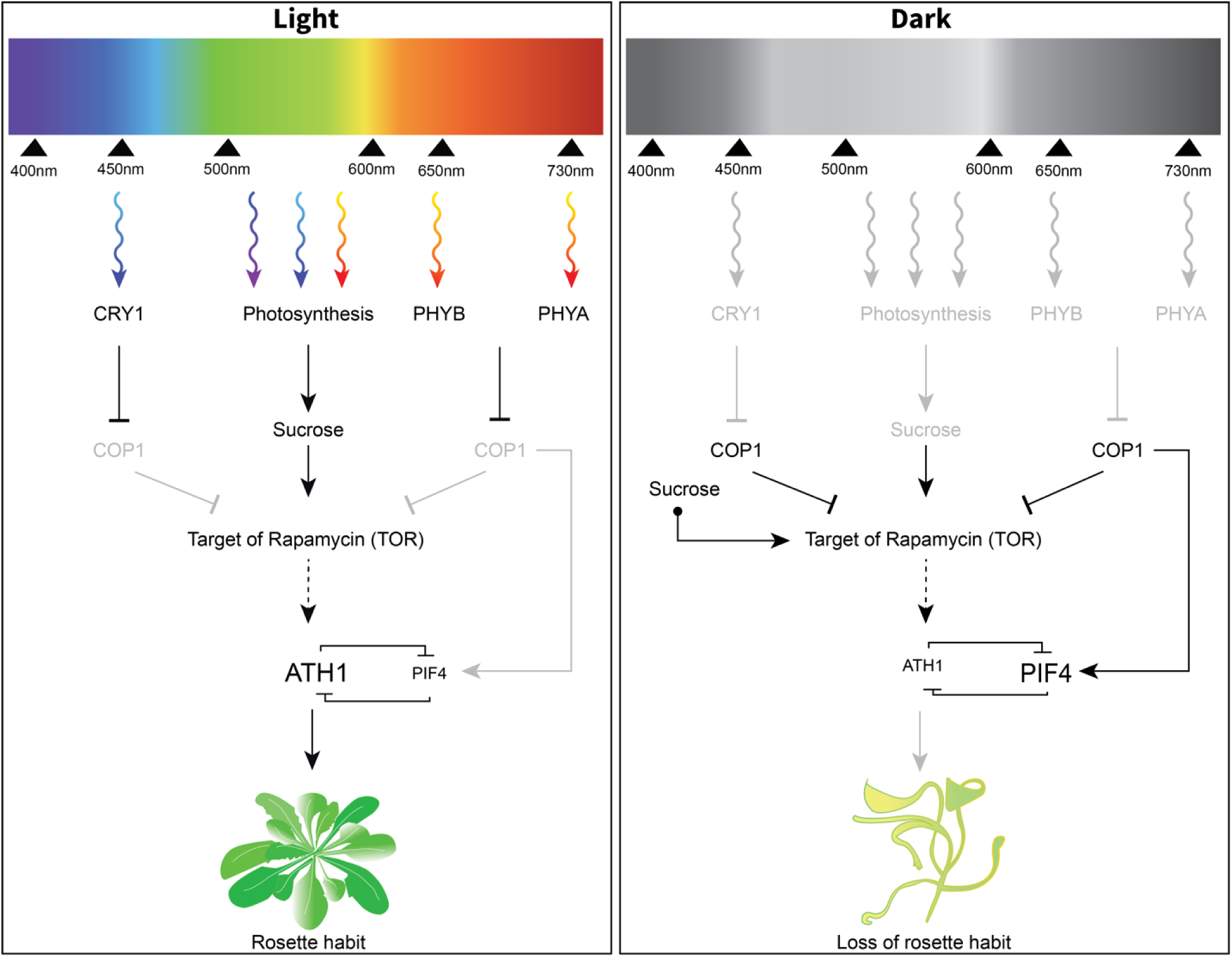
Light and sucrose signaling pathways converge at TOR kinase to control *ATH1* expression and subsequent rosette growth habit in *Arabidopsis thaliana*. Expression of *ATH1* is mediated by the activity of TOR kinase in response to both sugar and light. In response to light (**left panel**), photoreceptor signaling inhibits the activity of a COP1-containing protein complex that acts as a central repressor of light signaling in darkness. This releases the inhibitory effect of COP1 on TOR kinase. Activation of TOR kinase then leads to both activation of the SAM and induction of *ATH1* expression in the SAM. As a consequence of *ATH1* expression in the SAM, *PIF* gene expression, including *PIF4*, is locally inhibited. This contributes to inhibition of rib zone activity and, consequently, suppression of rosette internode elongation with the for Arabidopsis typical rosette growth habit as a result. As TOR kinase is a major regulator of mRNA translation, the effect on *ATH1* expression is most likely indirect (dotted arrow). In the absence of light (**right panel**), the COP1-complex is stabilized and inhibits TOR kinase activity and subsequent SAM activation. In addition, the COP1-complex stabilizes PIFs in darkness to positively regulate skotomorphogenesis. As a combined effect, *ATH1* is not expressed under these conditions. Sucrose-availability to the SAM can substitute for light both in the case of SAM activation and for *ATH1* induction. Although both processes are mediated through TOR kinase, sucrose levels sufficient to activate the SAM only result in weak expression of *ATH1*, probably as the result of still active COP1-PIF signaling. Resulting ATH1 levels are insufficient to suppress rib zone activity. As a consequence, in most circumstances sugar-induced dark-grown seedlings display a caulescent growth habit due to premature rib zone activation resulting in elongation of vegetative internodes.

## Discussion

In plants most of the adult body is formed post-embryonically by the continuous activity of pools of undifferentiated progenitor cells: the shoot and root apical meristems. Both apical meristems are established during plant embryogenesis. At the completion of embryogenesis the apical meristems are quiescent, but become reactivated after seed germination. In *A. thaliana* (Arabidopsis), light is crucial for reactivation of the shoot apical meristem (López-Juez et al., 2008; Arsovski et al., 2012; Pfeiffer et al., 2016; Mohammed et al., 2017). A direct outcome of light-activation of the meristem is the production of leaves. In rosette plants, such as Arabidopsis, the newly formed leaves give rise to a basal rosette: a whorl of leaves that has little to no elongation between successive nodes. The rosette habit is widespread amongst flowering plants and provides several advantages compared to taller, less compact plants, such as protection from (a)biotic stresses (Schaffer and Schaffer, 1979; Bello et al., 2005; Larcher et al., 2010; Thomson et al., 2011; Fujita and Koda, 2015). In Arabidopsis, light requirement for SAM activation can be overcome by availability of metabolizable sugars, such as sucrose, to the meristem (Roldán et al., 1999; Li et al., 2012). However, under such conditions plants fail to establish a typical compact rosette. Instead, they display a caulescent growth habit due to elongation of rosette internodes. Here we show that this dramatic change in growth habit in the absence of light is caused by premature activation of the rib zone due to insufficient expression of the light-mediated TALE homeobox gene *ATH1* in the shoot meristem. Our observations confirm a fundamental role for ATH1 in Arabidopsis rosette growth habit and supports a role for TOR kinase as a central hub for integration of energy and light signaling in controlling cell differentiation and organ initiation at the SAM. Previously, both activation of the SAM following germination, via induction of *WUSCHEL* (*WUS*), as well as subsequent initiation of leaf primordia were shown to be synergistically controlled by light-signaling pathways and photosynthesis-derived sugars, both conveyed by TOR kinase (Li et al., 2012; Pfeiffer et al., 2016). Here we show that induction of *ATH1* expression at the SAM, required to inhibit rib zone activation during vegetative development, is additively induced by sugar and light-signaling and that TOR kinase activity is essential for both inputs. Thus, TOR kinase integrates light and energy signals to activate the central population of stem cells in the SAM and subsequent differentiation processes at the periphery of the meristem but, interestingly, also represses differentiation processes by inhibiting rib zone activity at the basal part of the meristem. Induction of *ATH1* through these signals might indirectly result from SAM activation. However, we think this is unlikely as, in the absence of sucrose, the SAM of dark-grown *cop1* mutants remains dormant, while *ATH1* is expressed to relatively high levels. In addition, SAM activation and *ATH1* induction responses differ in their sensitivity to sucrose. Sucrose concentrations adequate to activate the SAM and initiate organogenesis, fail to induce significant levels of *ATH1*.

How TOR kinase controls *ATH1* promoter activity is currently unknown. TOR kinase is a major regulator of mRNA translation in all eukaryotic cells (Schepetilnikov and Ryabova, 2017). Moreover, light/dark transitions are also known to modulate mRNA translation in plants (Liu et al., 2012). Therefore, the effect of TOR kinase on *ATH1* is most likely indirect, as was recently reported for WUS by Janocha et al. (2021) who show that TOR kinase indirectly induces *WUS* expression by controlling cytokinin levels in the SAM through translational regulation of cytokinin catabolic enzymes.

In the absence of light, *ATH1* gene expression is repressed by negative regulators of photomorphogenesis, including COP1 (Quaedvlieg et al., 1995). This fits well with the observation that in darkness COP1 represses TOR kinase (Chen et al., 2018). Here we report that, in darkness, sucrose can substitute for light to induce *ATH1* expression and this also requires TOR kinase activity (Figure 7). As sucrose-mediated induction of ATH1 can still be observed in a *cop1* mutant background, sucrose affects TOR kinase activity most likely independently of COP1. Light signaling inactivates COP1, resulting in an induction of auxin biosynthesis. Auxin then activates the small Rho-like GTPase ROP2, which in turn activates TOR (Cai et al., 2017; Li et al., 2017; Schepetilnikov et al., 2017). Constitutive expression of activated ROP2 stimulates TOR in the shoot apex and is sufficient to promote organogenesis in the absence of light (Li et al., 2017). Sugars are known to trigger the accumulation of auxin, along with its biosynthetic precursors and this could be how sucrose activates TOR kinase in darkness, in the presence of COP1 (Chourey et al., 2010; LeClere et al., 2010; Sairanen et al., 2012; Mohammed et al., 2017). Worth mentioning in this respect is that the same set of PIF proteins that we identified as repressors of *ATH1* expression, negatively regulate sugar-induced auxin biosynthesis (Sairanen et al., 2012).

In the presence of light, expression of *ATH1* is induced in a functionally redundant manner by multiple cryptochrome and phytochrome photoreceptors operating in response to broad wavelengths from blue to far-red light (Figure 7). This ensures presence of *ATH1* in the SAM under all light conditions and thus inhibition of rib zone activity, resulting in the characteristic compact rosette growth habit of Arabidopsis plants. In line with this, loss of rosette habit has been observed in light-grown Arabidopsis plants lacking multiple functional phytochrome and/or cryptochrome photoreceptors. Control of vegetative internode elongation in response to changes in light quality or ambient temperature was shown to be mediated by the concerted action of phyA, phyB, phyD, phyE, and/or CRY1, all of which we identified here as having a role in light-mediated induction of *ATH1* expression (Devlin et al., 1996; Whitelam and Devlin, 1997; Devlin et al., 1998; Whitelam et al., 1998; Devlin et al., 1999; Mazzella et al., 2000; Devlin et al., 2003; Franklin et al., 2003a; Kanyuka et al., 2003; Strasser et al., 2010; Zhang et al., 2017). Often not appreciated in literature, compact rosette growth is thus a genuine photomorphogenic trait in Arabidopsis. Remarkably, rosette growth habit is a non-plastic trait, unlike other photoreceptor-driven developmental responses in Arabidopsis, such as elongation of hypocotyl, petiole, and inflorescence stem. A compact rosette habit is not affected in wildtype plants even under light quality conditions or temperature regimes that cause rapid elongation of aerial plant organs. Plasticity of growth and development is often considered adaptive, enabling sessile plants to adjust rapidly to a changing environment (Schlichting, 1986; Schlichting and Levin, 1986). However, as mentioned, rosette growth habit provides several advantages compared to a caulescent growth habit. Loss of a compact rosette in response to environmental cues, therefore, might be detrimental to plant fitness and viability. A compact rosette habit is not constitutively expressed in all rosette species (our unpublished observations) and this trait, as a result of selection, may have become fixed in Arabidopsis through genetic assimilation (Ehrenreich and Pfennig, 2016). Important contributors to genetic assimilation are genetic variants that alter gene regulation. Plausible ways in which gene regulation might facilitate loss of phenotypic plasticity are i) decoupling of the regulation of genes that control a plastic trait from environmental cues or ii) the evolution of additional regulatory pathways that makes their expression insensitive to the environment (Ehrenreich and Pfennig, 2016). The latter explanation (ii) might be true in the case of Arabidopsis rosette growth habit, given that *ATH1* expression is induced in response to broad wavelengths and involves multiple photoreceptors. Moreover, it has been proposed that ATH1 controls internode elongation by antagonizing a large number of genes that promote internode growth, mostly independent of each other (Ejaz et al., 2021). This assumption fits with the observation that *pifq* reduced internode elongation in *ath1-3* only by about 70%. Such multitarget control by ATH1 of genes that affect internode elongation would further contribute to the robustness of rosette growth habit in *A. thaliana*. Therefore, it is of interest to investigate whether ATH1 has a similar role in other rosette species and, if so, whether differences in plasticity of rosette compactness can be linked to differences in light-signaling control of *ATH1* expression and/or decoupling genes that affect internode elongation from ATH1 regulation.

In this study, we identified *PIF1*, *PIF3*, *PIF4*, and *PIF5* as transcriptional targets of ATH1. Of these, *PIF4* and several steps of the PIF pathway, were previously identified as a binding target of ATH1 (Ejaz et al., 2021). Therefore, ATH1 might affect the expression of these four *PIF* genes through direct transcriptional repression. Our finding that PIF4, and at least one of the other PIF proteins, PIF1, PIF3, or PIF5, in turn function as negative regulators of *ATH1* expression suggest the presence of a double-negative feedback loop between ATH1 and PIF family members (Figure 7). Signaling systems like ATH1 and PIFs that contain double-negative feedback loops can, in principle, convert graded inputs into switch-like, irreversible responses (Ferrell, 2002). Such a genetic toggle switch is a bistable dynamical system, possessing two stable equilibria, each associated to a fully expressed protein. ATH1 has a fundamental role in maintaining the Arabidopsis rosette habit during vegetative growth. When grown in the presence of light, *ATH1* is expressed throughout the shoot meristem, including the subapical region where it represses stem growth. Plant switching to reproductive growth rapidly downregulate *ATH1* at the shoot meristem, marking the onset of bolting and emergence of an elongated inflorescence (Proveniers et al., 2007; Gómez-Mena and Sablowski, 2008; Ejaz et al., 2021). Such stem elongation is absent in plants constitutively expressing *ATH1,* without affecting formation of flowers (Cole et al., 2006; Gómez-Mena and Sablowski, 2008; Rutjens et al., 2009). The data presented in this study show that both absence of ATH1 and induced ATH1 expression, leads to pronounced changes in *PIF* gene expression at the SAM associated with significant elongation or complete suppression of rosette internodes, respectively. PIF proteins have been associated with bolting time and/or stem internode elongation (Brock et al., 2010; Todaka et al., 2012; Galvāo et al., 2019; Arya et al., 2021; Jenkitkonchai et al., 2021). Moreover, elongated rosette internodes can be observed *35S::PIF4* plants (see Figure 1d in Kumar et al. (2012)). It is therefore proposed that an ATH1-PIF toggle switch underlies the rapid and distinctive switch in Arabidopsis growth habit that marks floral transition.

## Methods

### Plant material and growth conditions

Arabidopsis seeds were obtained from the Nottingham Arabidopsis stock center (http://arabidopsis.info/; stock number between brackets) or were kind gifts of colleagues. The following genotypes were used: Col-8 wild type (N60000), L*er* wild-type (NW20), *phyB-5* (Reed et al., 1993), *phyBDE* (Shalitin et al., 2002), *phyABDE* (Franklin et al., 2003b), *phyBcry1* (Mazzella et al., 2000) *cry1-1* (Ahmad and Cashmore, 1993), *hy1-1* (Muramoto et al., 1999), *hy1cry1*, *hy1cry2*, *hy1cry1cry2* (López-Juez et al., 2008), *pif3-1* (N530753; Kim et al., 2003), *pif4- 1* (N667486; Huai et al., 2018), *pif4pif5* (N68096; Leivar et al., 2012)*, pif7-1* (N68809), *pif3pif4* (N66046), *pif1pif3* (N66045), *pif1pif3pif4* (N66500), *pif1pif3pif5* (N66047), *pif3pif4pif5* (N66048), *pif1pif3pif4pif5* (*pifq*; N66049) (Leivar et al., 2008), *cop1-4* (McNellis et al., 1994), *Pro_35S_:HA-ATH1*, *ATH1pro:GUS*, *ath1-3* (Proveniers et al., 2007), *ath1-4* (Li et al., 2012) and *35Spro:ATH1-HBD* lines (Rutjens et al., 2009), *35S::ALCR alcA:RNAi-TOR* and *35S:ALCR alcA:GUS* (Deprost et al., 2007).

To introduce *ATH1pro:GUS* in Col-8, L*er*, *phyB*, *phyBcry1*, and *phyBDE* backgrounds, the original *ATH1pro:GUS* line was crossed into the respective genotypes and the resulting offspring was backcrossed at least four times to the parental acceptor lines. For ectopic expression of *ATH1* in photoreceptor mutants, L*er*, *phyB*, *phyBcry1*, *phyBDE*, and *hy1cry1cry2* plants were transformed with a *HA*-tagged version of *ATH1* (*Pro_35S_:HA-ATH1*) as described previously (Proveniers et al., 2007). Transgenic seeds were selected based on seed GFP expression. Per genotype over ten independent, homozygous single insert lines were used for further analysis. To obtain *ath1-3 pifq* plants, F2 offspring plants from a cross *ath1-3* x *pifq* were preselected based on a, for higher-order *pif* mutants, characteristic short-petiole phenotype. Selected plants were then genotyped for homozygous presence of all five mutant alleles using the primers listed in Supplemental Table S2. To obtain *cop1-4 ath1-4* plants, F2 offspring plants from a cross *cop1-4* x *ath1-4*, homozygous for the *cop1* mutation, were selected under dark conditions for a short-hypocotyl phenotype. Subsequently, selected seedlings were transferred to normal growth conditions and genotyped for the *ath1-4* mutation.

For plant growth, seeds were chlorine gas sterilized in a desiccator for 4 h using a mixture of 4 ml 37% HCl and 100 ml commercial bleach (Glorix; contains 4.5% active chlorine) and put on soil (Primasta) or on sterile 0.8% plant agar (Duchefa Biochemie) with full strength Murashige-Skoog medium (MS salts including MES Buffer (pH 5.8) and vitamins, Duchefa Biochemie) in square Petri dishes (120x120 mm). After 2-3 days of stratification (4°C, darkness), plants were grown in climate-controlled growth cabinets (Snijders, Microclima 1000) in either a short day (SD; 8 hours light/16 hours dark) or a long day (LD; 16 hours light/8 hours dark) photoperiod, under 120 µmol m^−2^ s^−1^ photosynthetic active radiation (PAR) fluorescent white-light conditions (Sylvania, Luxline Plus Cool White) and 70% relative humidity. For growth of plants under monochromatic light conditions, plants were grown in a Snijders Microclima growth cabinet equipped with Philips GreenPower LEDs (red light (124.35 µmol m−2 s−1), blue light (6.14 µmol m−2 s−1) or far-red light (77.57 µmol m−2 s−1)).

For liquid culture, Arabidopsis seedlings were grown in 100 mL bottles on a rotary shaker (185 rpm, 22 °C). Per bottle, ten to twenty seeds were added to 20 mL half-strength MS medium (MS salts including MES Buffer (pH 5.8) and vitamins, Duchefa Biochemie) and bottles were sealed with Steristoppers® (Heinz Herenz, Hamburg). After stratification, seeds were exposed to fluorescent light (1-1.5 hours, 120 µmol/m2/s) to stimulate germination, after which the bottles were wrapped in aluminum foil to avoid any further light exposure. Unless stated otherwise, sterile sucrose (50% w/v) or sorbitol (50% w/v) was added to a final concentration of 1% v/v either at the beginning or at day three of the experiment.

### Phenotypic analyses

#### Rosette internode elongation

For light-grown plants, total rosette internode length was measured using a digital caliper. For dark-grown seedlings, plants were photographed after three weeks of growth in liquid medium and ImageJ (Schneider et al., 2012) was used to measure total rosette internode length. To determine the average rosette internode length total rosette internode length was divided by the total number of rosette leaves. In light-grown plants, the first leaf to give rise to a secondary inflorescence was defined as the first cauline leaf. Dark-grown plants were never grown long enough to reach the reproductive phase.

#### Meristem cell size

Using confocal laser scanning microscopy, longitudinal optical sections were made through apices of five-day-old seedlings. In median sections a central cell file extending from the epidermis into the subapical region where the hypocotyl vascular strands converge was identified. Using ImageJ software ((NIH; https:imagej.nih.gov/ij/), individual cell lengths were then measured per position in apical-basal direction.

### Gene expression analysis (qPCR)

All samples (whole seedlings or SAM-enriched tissue) were snap-frozen in liquid N2 and stored at -80°C before RNA extraction. For each experiment 3 or 4 biological replicates and two technical replicates were included. RNA was isolated using a RNeasy mini or micro kit (Qiagen, http://www.qiagen.com/). Genomic DNA was removed using DNaseI (Thermo Scientific) and cDNA was synthesized from 500 ng - 1 µg RNA using RevertAid H Minus Reverse Transcriptase and Ribolock RNAse inhibitor (Thermo Scientific) using a mix of anchored odT(20) primers (Jena Bioscience) and random hexamers (IDT). qPCR reactions were performed using qPCRBIO SyGreen Blue mix (PCRBIO) on a ViiA7 Real Time PCR system and ViiA7 software was used to analyze the data. Relative expression levels were calculated using the ΔΔCt method (Livak and Schmittgen, 2001), normalized to the expression of the reference genes *GAPC2* (AT1G13440) and/or *MUSE3* (AT5G15400). Differences between mutant and wild-type ΔΔCt values were statistically analyzed with an independent sample t-test or ANOVA test (p < 0.05) in Rstudio version 1.2.5033. For primer sequences used, see Supplemental Table S1.

### β-Glucuronidase Staining and Microscopy

Seven-day-old seedlings were harvested and vacuum-infiltrated in β-glucuronidase (GUS) staining buffer (50 mM sodium phosphate buffer (pH=7.2), supplemented with 0.1% Triton X-100, 100 mM, potassium ferrocyanide, 100 mM, potassium ferricyanide, 2 mM 5-bromo-4-chloro-3-indolyl glucuronide). Samples were incubated at room temperature for 16 hours and subsequently cleared in ethanol baths with increasing concentration (20%, 30%, 50%, and 70%). Images were taken with a Nikon DXMI200 camera attached to a Zeiss Stemi SV11 stereo microscope. The GUS staining area in the shoot apex was measured and quantified using Image J (NIH; https://imagej.nih.gov/ij/).

### Confocal microscopy

Five-day-old Col-8, *ath1-4*, and *35Spro:ATH1-HBD* seedlings were grown in either continuous darkness supplied with one percent sucrose or in continuous light at 27°C. Seedlings were cleared using the ClearSee method (Kurihara et al., 2015) and imaged at a resolution of 0.25 x 0.25 x 0.5 µm using a Confocal Laser Scanning microscope Carl Zeiss LSM880 Fast AiryScan with a Plan-Apochromat 63x/1,2 Imm Korr DIC objective (numerical aperture 1.40, oil immersion) and ZEN software (blue edition, Carl Zeiss). Staining of apices with Calcofluor White Stain (Sigma-Aldrich) was performed as described before (Ursache et al., 2018). Excitation was at 405 nm and emission filters were set between 425 nm and 475 nm. All replicate images were acquired using identical confocal microscopy parameters for each experiment. Confocal images were processed with Fiji (version 1.52, Fiji) and Adobe Illustrator.

### Statistical analysis

Data plotting and statistical analysis were performed using RSTUDIO.1.0.143 (www.rstudio.com) with R v.3.6.2 (https://cran.r-project.org/). The significance of differences between experimental groups was determined by using the student’s t-test as indicated in each experiment. The significance level of differences between two experimental groups was either marked as *, p<0.05; **, p<0.01; ***, p<0.001; ns, not significant or indicated by the corresponding P values. In order to compare the significance of differences between multiple experimental groups, the Fisher’s least significant difference (LSD) test and one-way analysis of variance (ANOVA) were performed using the Bonferroni correction with an α = 0.05 from the agricolae package (Mendiburu, 2020). Images containing micrographs and other images were compiled using Adobe Illustrator and ImageJ software (https://imagej.nih.gov/ij/) to create the figures.

## Author contributions

Conceptualization: M.P.; Methodology: M.P., S.S.H., S.S.S., and E.S.; Investigation: S.S.H., E.S., N.P., G.B., and S.S.S.; Writing – Original Draft: M.P. and S.S.H.; Writing – Review & Editing, M.P., S.S.H., S.S.S., E.S. and S.S.; Funding Acquisition: M.P., S.S.H., S.S. ; Visualization: S.S.H.; Resources: M.P., E.S., and S.S.; Supervision: M.P. and S.S.

## Acknowledgements

*phyA cry1*, *phyB cry1* and *phyA phyB cry1* seeds were a kind gift of Jorge Casal. *hy1, cry1, cry2, cry1 cry2, hy1 cry1, hy1 cry2, and hy1 cry1 cry2* seeds were a kind gift of Enrique Lopez-Juez and *cop1-4* seeds were kind gift of Jan Lohmann.

---

**Figure S1:**
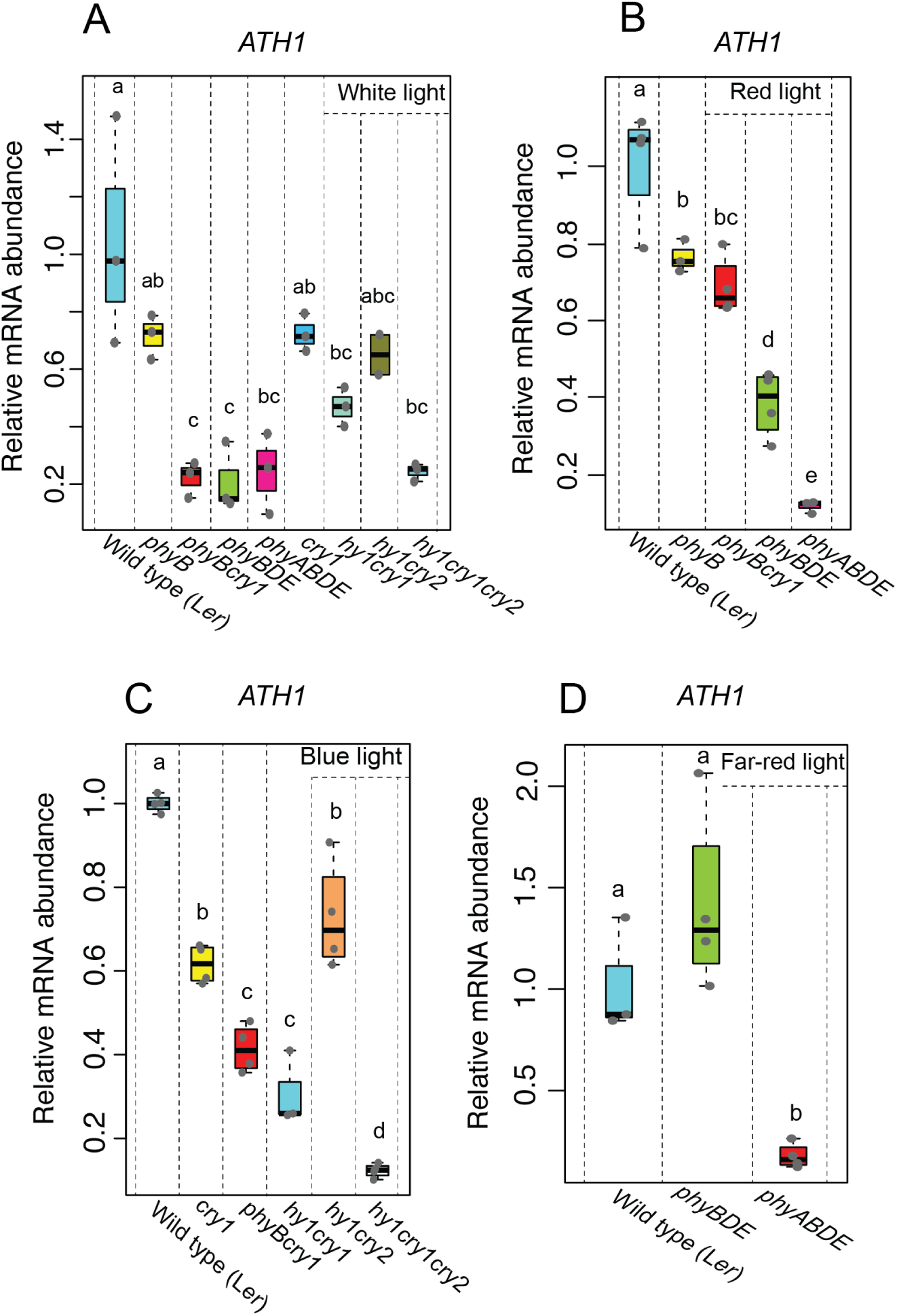
Relative mRNA abundance of ATH1 in different photoreceptor mutants. **A-D**: Relative expression of *ATH1* in seven-day-old wild type (L*er*) and indicated photoreceptor mutants grown under SD conditions in the presence of white light (**A**), red light (**B**), blue light (**C**) or far-red light (**D**). Transcript levels were normalized to *GAPC2* (AT1G13440) (BL) or *MUSE3* (AT5G15400; WL, RL, and FRL). Data shown are the average of three biological replicates. At least 30 seedlings were used for each biological replicate. Different letters denote statistically significant differences between groups (P < 0.05) as determined by 1-way ANOVA followed by Tukey’s post hoc test.

**Figure S2:**
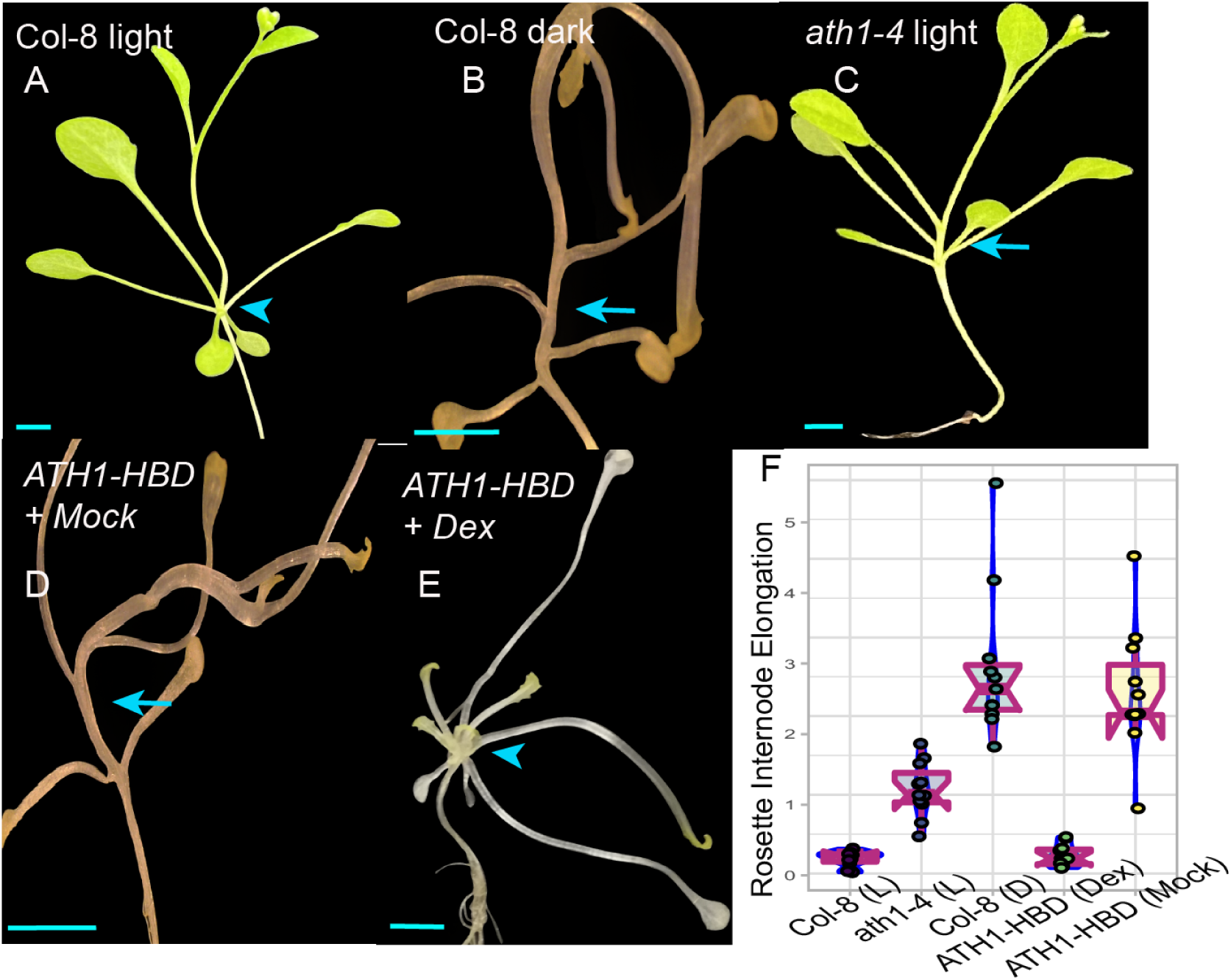
Sugar-induced dark-grown seedlings phenocopy light-grown *ath1*mutants. Rosette elongation phenotypes of wild-type Col-8 (**A** and **B**) *ath1-4* (**C**), and *35Spro:ATH1-HBD* (*ATH1-HBD*; **D** and **E**) plants grown in the presence (**A**, **C**) or absence (**B**, **D**, **E**) of light at 27°C. *35Spro:ATH1-HBD* plants were treated either with a mock (0.1% ethanol, **D**) or 10 μM dexamethasone (Dex, **E**). Dark-grown plants (**B**, **D**, **E**) were supplemented with sucrose to a final concentration of one percent three days after the start of the experiment. Arrows indicate elongated rosette internodes, the arrowhead indicates complete suppression of internode elongation. **F:** Average rosette internode lengths of plants depicted in **A**-**E**. Per genotype and treatment 10 individual plants were analyzed. Scale bars represent 5 mm.

**Figure S3:**
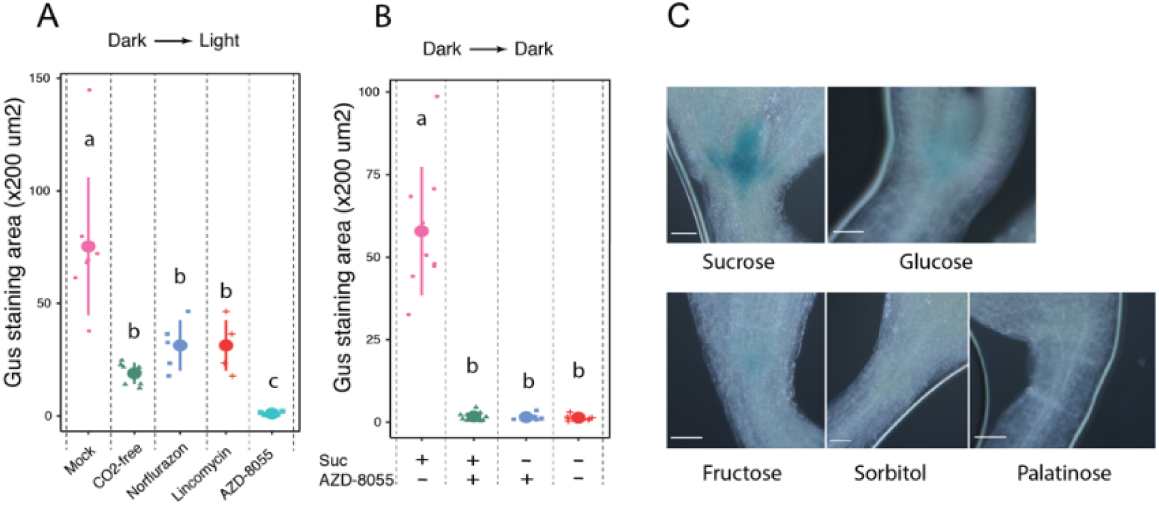
*ATH1_pro_:GUS* activity in seven-day-old seedlings grown in darkness in the presence of different sugars. **A, B**: Quantification of GUS-staining intensity in *ATH1_pro_:GUS* shoot apices from Figure 5B, E. Different letters denote statistically significant differences between groups (P < 0.05) as determined by 1-way ANOVA followed by Tukey’s post hoc test. **C**: Shoot apices of GUS-stained *ATH1_pro_:GUS* seedlings grown in continuous darkness for seven days. Plants were grown in the presence of either sucrose, glucose, fructose, palatinose, or sorbitol, all added to the growth medium to a final concentration of one percent at the start of the experiment. Scale bars represent 0.05 mm.

**Figure S4:**
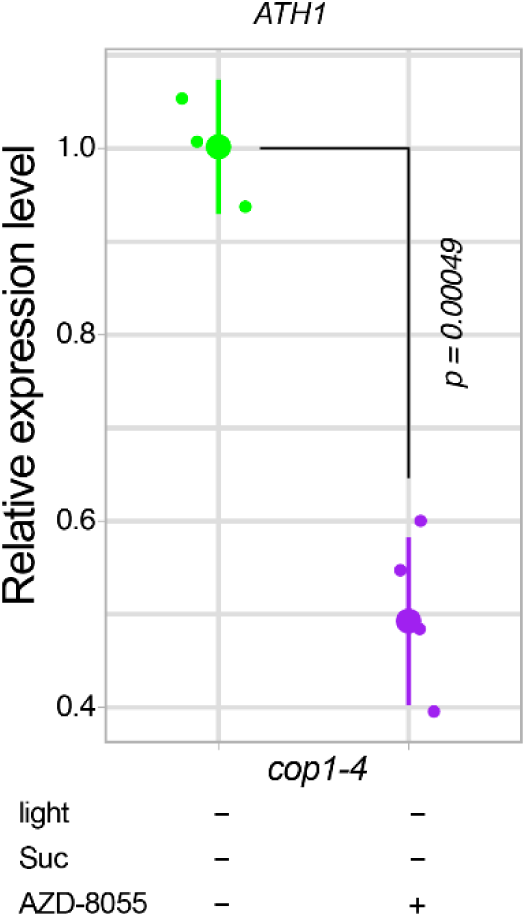
Effect of TOR inhibition on *ATH1* expression in *cop1-4* mutant seedlings. Relative expression of *ATH1* in seven-day-old *cop1-4* mutants grown according to the experimental setup indicated in Figure 5A. Seeds were light-treated for 45 minutes to stimulate germination and then grown in continuous darkness for five days. Following the addition of AZD- 8055, the seedlings were grown for two more days in darkness before samples were taken. Transcript levels were normalized to *MUSE3* (AT5G15400). The average of three (AZD-8055 -) or four (AZD-8055 +) biological replicates is shown. At least 30 seedlings were used for each biological replicate. The p-value of significant difference, as determined by the two-tailed Student’s t-test, is indicated in the figure.

**Supplemental Table S1:**
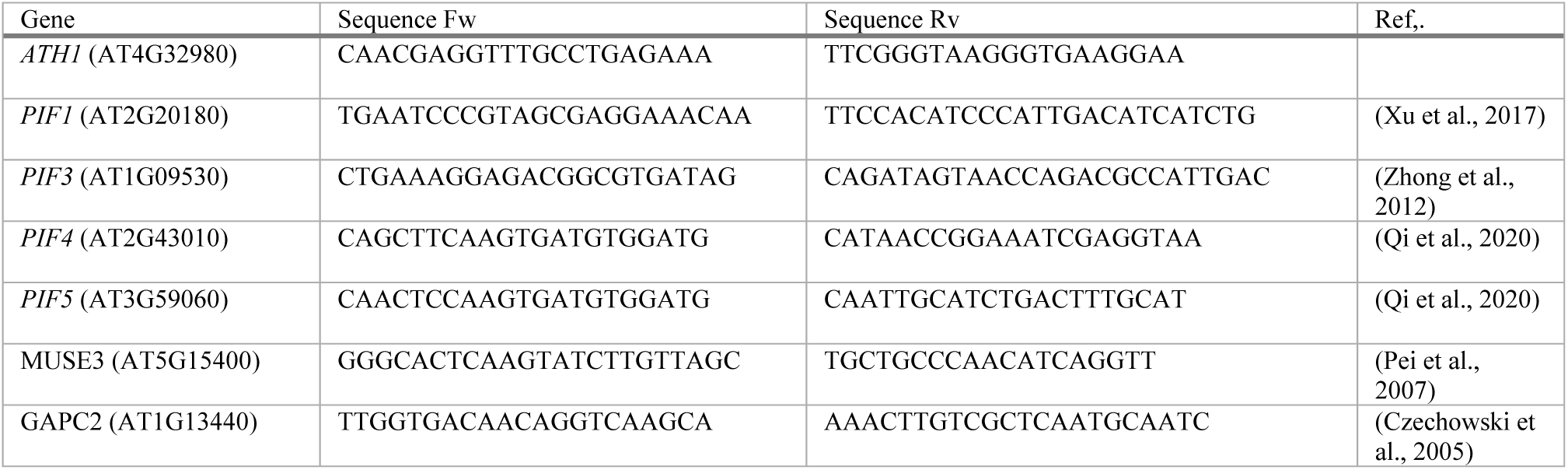
Primers for qRT-PCR.

**Supplemental Table S2:**
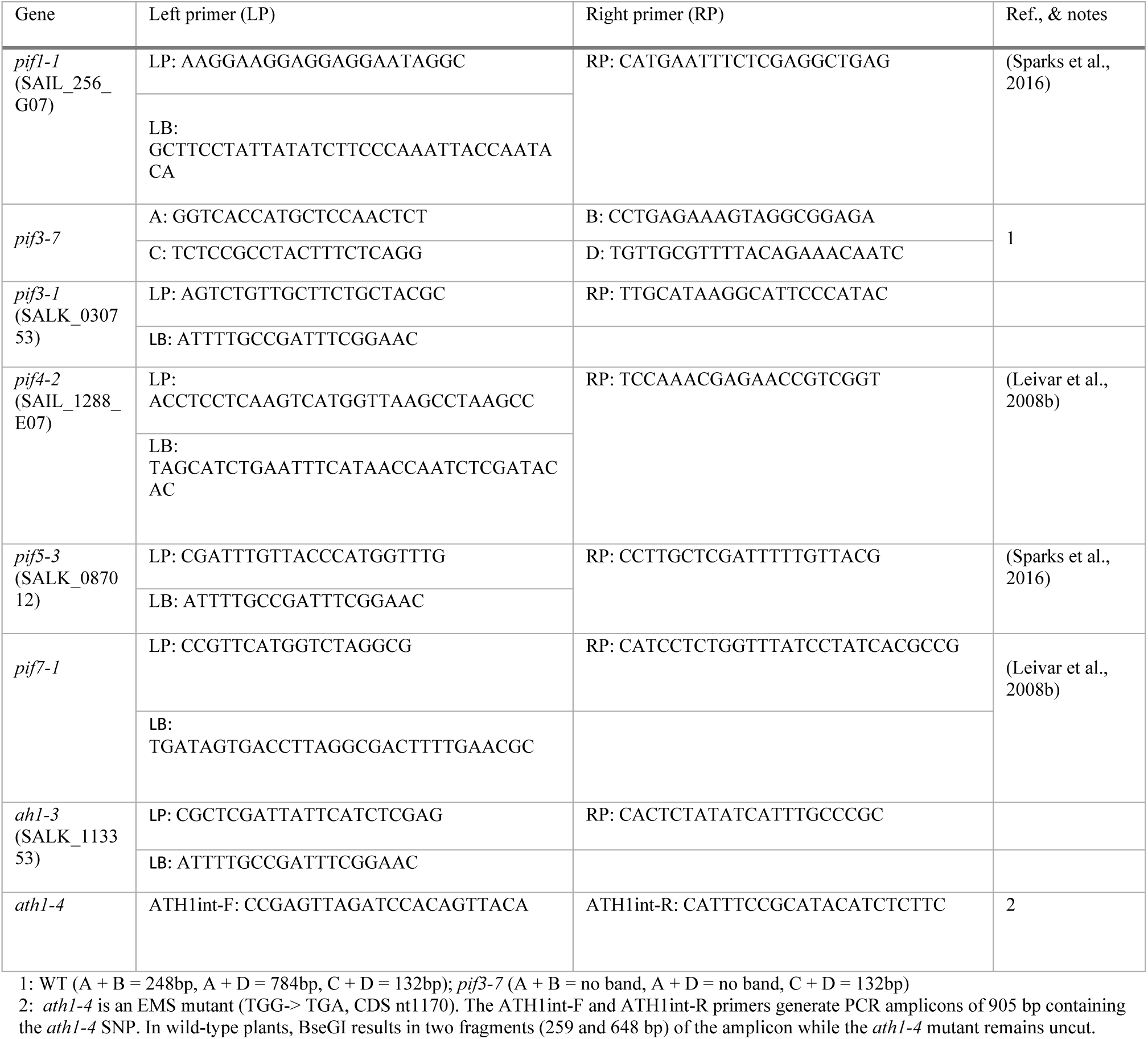
Primers for genotyping.

## Notes

### Competing Interest Statement

The authors have declared no competing interest.

